# Emergence and fragmentation of the alpha-band driven by neuronal network dynamics

**DOI:** 10.1101/2021.07.19.452820

**Authors:** Lou Zonca, David Holcman

**Affiliations:** Sorbonne University, Pierre et Marie Curie Campus, 5 place Jussieu 75005 Paris, France; Group of Applied Mathematics and Computational Biology, École Normale Supérieure, France

## Abstract

Rhythmic neuronal network activity underlies brain oscillations. To investigate how connected neuronal networks contribute to the emergence of the *α*-band and the regulation of Up and Down states, we study a model based on synaptic short-term depression-facilitation with afterhyperpolarization (AHP). We found that the *α*-band is generated by the network behavior near the attractor of the Up-state. Coupling inhibitory and excitatory networks by reciprocal connections leads to the emergence of a stable *α*-band during the Up states, as reflected in the spectrogram. To better characterize the emergence and stability of thalamocortical oscillations containing *α* and *δ* rhythms during anesthesia, we model the interaction of two excitatory with one inhibitory networks, showing that this minimal network topology leads to a persistent *α*-band in the neuronal voltage characterized by dominant Up over Down states. Finally, we show that the emergence of the *α*-band appears when external inputs are suppressed, while the fragmentation occurs at small synaptic noise or with increasing inhibitory inputs. To conclude, interaction between excitatory neuronal networks with and without AHP seems to be a general principle underlying changes in network oscillations that could apply to other rhythms.

**Author summary:** Brain oscillations recorded from electroencephalograms characterize behaviors such as sleep, wakefulness, brain evoked responses, coma or anesthesia. The underlying rhythms for these oscillations are associated at a neuronal population level to fluctuations of the membrane potential between Up (depolarized) and Down (hyperpolarized) states. During anesthesia with propofol, a dominant alpha-band (8-12Hz) can emerge or disappear, but the underlying mechanisms remain unclear. Using modeling, we report that the alpha-band appears during Up states in neuronal populations driven by short-term synaptic plasticity and noise. Moreover, we show that three connected networks representing the thalamocortical loop reproduce the dynamics of the alpha-band, which emerges following the arrest of excitatory stimulations, but can disappear by increasing inhibitory inputs. To conclude, short-term plasticity in well connected neuronal networks can explain the emergence and fragmentation of the alpha-band.

## Introduction

Electroencephalogram (EEG) is used to monitor the brain activity in various conditions such as sleep [1,2], coma [3] or meditation [4] and to reveal and quantify the presence of multiple frequency oscillations [5] over time [6]. This analysis can be used to asses the level of consciousness or depth of unconsciousness of the brain. For example, during general anesthesia under propofol, a dominant oscillation is the *α*-band (8-12Hz) [7,8]. However, the precise mechanisms underlying the emergence or disappearance of this *α*-band remain unknown. Interestingly, when the level of sedation becomes too high, the EEG shows that the *α*-band can get fragmented and even disappear replaced by a different transient motif called burst-suppression, which consists in alternation of periods of high frequency activity followed by iso-electric suppression periods where the EEG is almost flat [7]. In general, large doses of hypnotic in prolonged anesthesia in rodents alters brain synaptic architecture [9], confirming the need to avoid over sedation. Burst-suppression is a motif associated with a too deep anesthesia and its presence could indicate possible post anesthetic complications, although it has been attributed to ATP depletion [10]. Recently, it was shown that the loss of the *α*-band announces the appearance of burst-suppressions [11], however, this causality between *α*-band suppression and burst-suppressions remains unexplained.

The *α*-band revealed by the EEG signal is associated with the local Up and Down states activity [12–14], which corresponds to a depolarized and hyperpolarized membrane voltage of a neuron [15]. The alternation between Up and Down states generates slow wave oscillations present in NREM sleep, as reported in slices electrophysiology [16] as well as using modeling approaches [17,18]. Similarly, the emergence of the *α*-band during anesthesia could result from network interactions, as proposed by models based on the Hodgkin-Huxley formalism [19–21].

Since Up and Down states reflect the neuronal activity at the population level [15,22], we propose here to investigate the emergence and fragmentation of the *α*-band using a modeling approach based on synaptic short-term plasticity [23,24], which is often used to obtain estimations for burst or interburst durations [25–27]. These models based on facilitation and depression have recently been used to evaluate the working memory capacity to remember a sequence of words [28].

Here, we use a mean-field neuronal model that accounts for both synaptic short-term dynamics and afterhyperpolarization (AHP) [29] resulting in a refractory period during which neurons stops firing after a burst. As a result, at a population level, AHP can modify the type of oscillations [30], from waxing and waning spindle oscillations to slow waves.

We first study a single, two and then three interacting neuronal networks, a minimal configuration revealing the coexistence of *α*-oscillations and switching between Up and Down states. As we shall see, only the neuronal population with AHP can trigger spontaneous switching between Up and Down states while the other one, without AHP is at the origin of the *α*-oscillations in the Up state. We also investigate the role of synaptic noise and model the effect of propofol as an excitatory current for inhibitory neurons.

## 1 Results

### 1.1 EEG reveals the dynamics of the *α*-band during general anesthesia

General anesthesia can be monitored using EEG (fig. 1A) that often reveals a stable *α*-band which persists in time (fig. 1B). The origin of the *α*-band is not fully understood but it was found to result from the dynamics of neuronal populations involving the reciprocal connexions between the thalamus and the cortex. During anesthesia involving the propofol agent, the inhibitory neurons are activated resulting in the emergence and stability of the *α*-band. Increase of the anesthetic agent can lead to a deeper anesthesia characterized by a transient disapearance of the *α*-band (fig. 1C) so that the spectrum is carried by the *δ*-band. This disappearance can be quantified by two parameters, defining a fragmentation level which accounts for the persistence *P_α_* of the *α*-power over a specific period of time and the number of disruptions *D_α_* in the power band (see Methods section 2.1). The exact mechanisms leading to the stability and the peak of the *α*-band frequency remains unclear. In the remaining part of this manuscript we propose to develop mean-field models based on synaptic properties to address this question.

**Figure 1:**
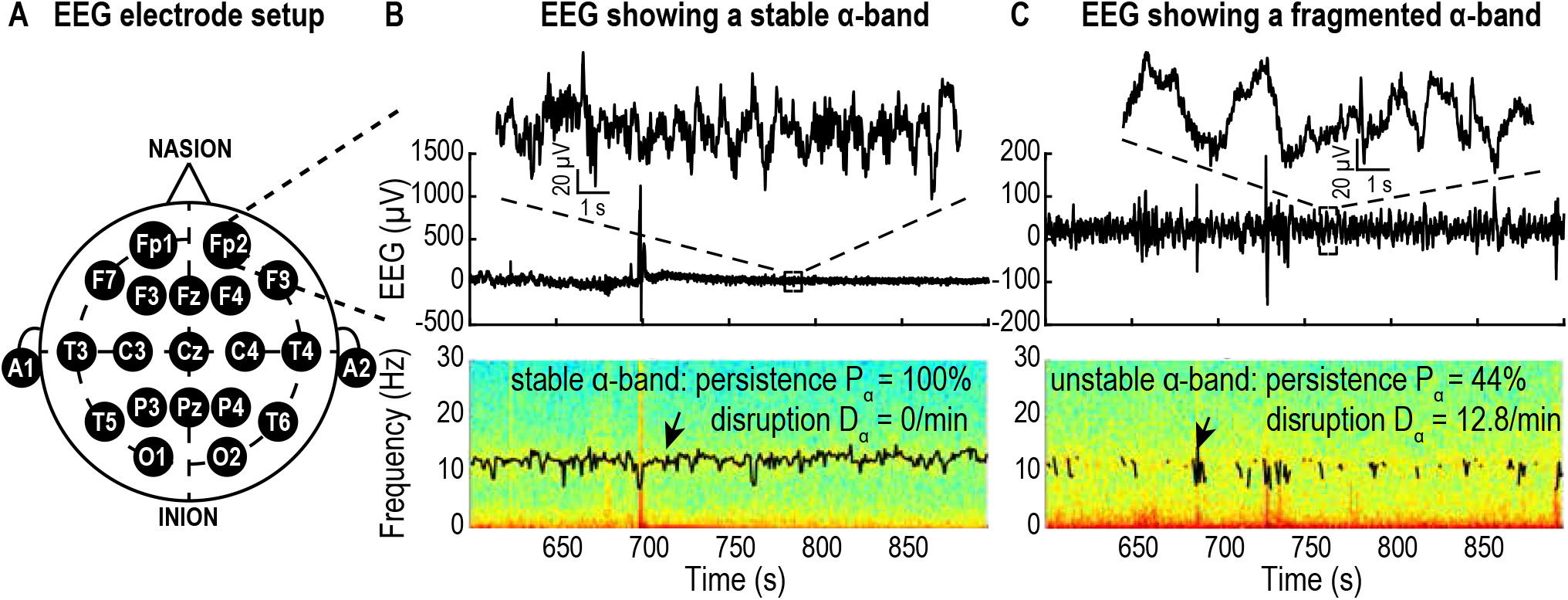
EEG recorded during general anesthesia. **A.** Schematic of EEG electrode setup on a patient’s head. **B.** Upper: Time recordings showing the EEG (inset: 5 s). The EEG signal is composed of multiple bands as shown in the spectrogram (lower panel) composed of two major bands: the *δ*-band (0-4Hz) and the *α*-band (8-12Hz) tracked by its maximum (black curve) revealing the persistence of the *α*-band during anesthesia. **C.** Same as A for a case of fragmented *α*-band. Data from the database VitalDB [31].

### 1.2 A single neuronal population can exhibit *α*-oscillations or slow waves through switching between Up and Down states

#### 1.2.1 The synaptic depression-facilitation model generates locked *α*-oscillations

To analyze the change between a persistent *α*-band and a *δ*-band we develop a mean-field model of neuronal networks based on short term synaptic plasticity (fig. 2A). The first model consists of one well connected population of excitatory neurons described by three variables: the mean voltage *h*, the synaptic facilitation *x* and the depression *y*, resulting in a stochastic dynamical system (see Methods, section 2.2, equations 3) showing bi-stability: one attractor corresponds to the Down state (hyperpolarized, low frequency oscillations) and the second one to the Up state (depolarized, high frequency oscillations). One fundamental parameter is the level of connectivity *J* that we shall vary (fig. 2B). We found that such a system can generate a dominant oscillatory band where the peak value is an increasing function of the connectivity *J* (fig. 2C). In the present scenario the network dynamics is locked into an Up state and the dominant oscillations are generated by the imaginary part of the eigenvalues at the Up state attractor. This result shows that the persistent oscillations are the consequence of the noise and of the synaptic properties as well as the biophysical parameters (Table 1, SI), indeed, changing the synaptic properties can lead to the fragmentation and disappearance of the band where most of the energy is now located in the *δ*-band as quantified by the spectral edge frequency at 95% (SEF95, fig. S1). In addition, varying the noise amplitude allows to either fragment the band (fig. 2D, *σ* = 7) or to increase the power and the persistence of the band (fig. 2D, *σ* = 15) but it does not affect the value of the peak of the dominant oscillation (fig. 2E). Indeed, similarly to the EEG analysis (fig. 1), we quantified the fragmentation level for different noise amplitudes and found that it varies from (*P_α_, D_α_*) = (52%, 34/*min*) for *σ* = 5 to (86%, 16/*min*) for *σ* = 7 and to (100%, 0/*min*) for *σ* = 15, which is of the same order of magnitude as the fragmentation levels observed in the human EEG data. This fragmentation of the *α*-band results only from the changes in the noise amplitude and can occur even though the population is locked into the Up state, suggesting that the loss the *α*-band could be independent from the switch between Up and Down states.

**Figure 2:**
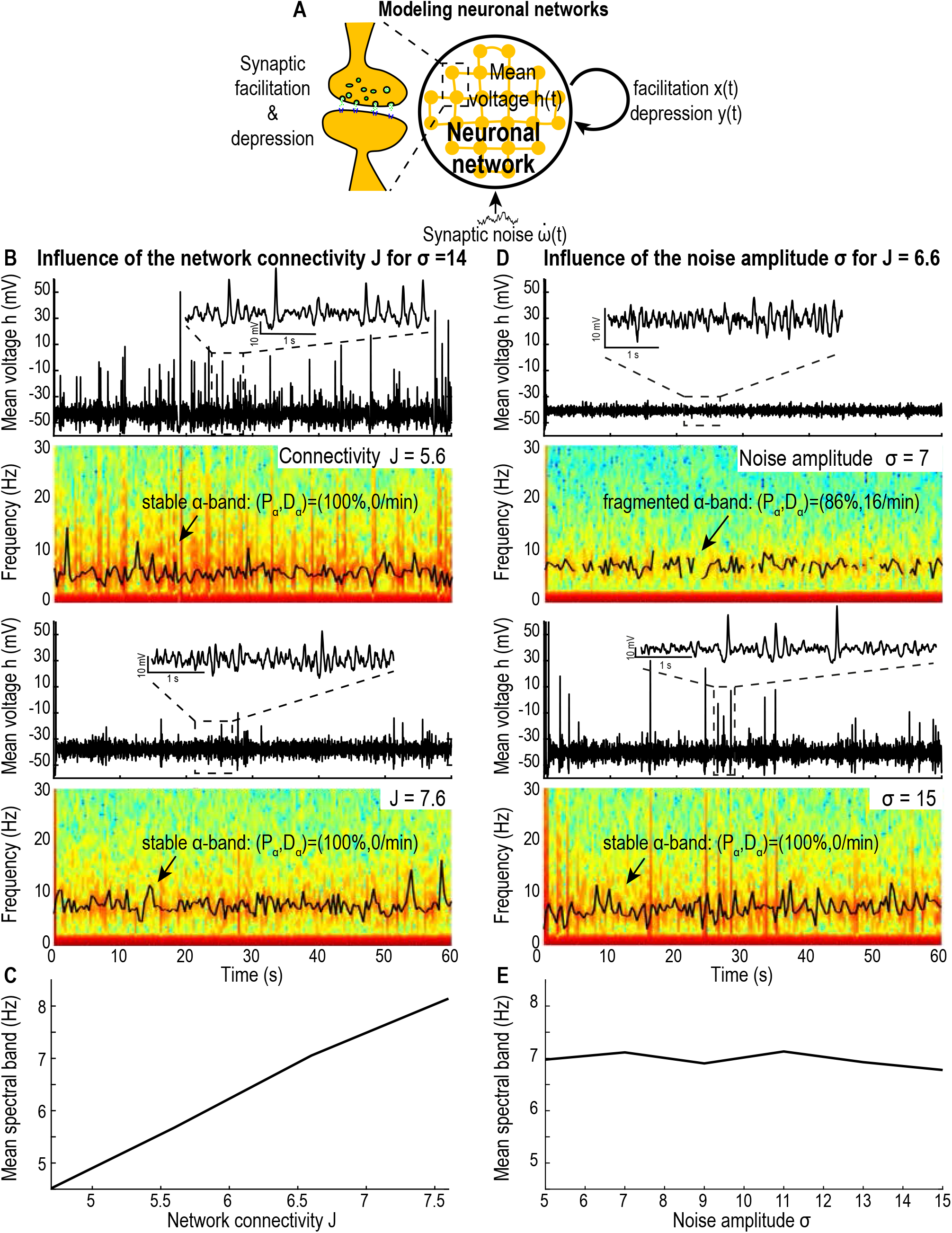
Effect of network connectivity *J* and noise amplitude *σ* on model (3) without AHP. **A.** Schematic of the facilitation-depression model (3). **B.** Time-series and spectrograms of *h* (60s simulations) with peak value of the dominant oscillatory band (black curve) for *J* = 5.6 (upper) and 7.6 (lower). **C.** Mean value of the dominant oscillatory frequency for *J* ∈ [4.8, 7.8]. **D.** Time-series and spectrograms of *h* (60s simulations) peak value of the dominant oscillatory band (black curve) for *σ* = 7 (upper) and 15 (lower). **E.** Mean value of the dominant oscillatory frequency for *σ* ∈ [5,15]. Synaptic plasticity timescales: *τ* = 0.01s, *τ_r_* = 0.2s and *τ_f_* = 0.12*s*.

#### 1.2.2 The synaptic depression-facilitation model with AHP generates Up and Down states but no *α*-oscillations

The previous model did not allow a dynamical switch between Up and Down states, thus we decided to test the effect of AHP in our model to explore a larger range of neuronal dynamics (see Methods). The synaptic depression-facilitation model with AHP is constructed by adding the AHP components to the mean-field equations (3) presented above (see Methods). The dynamics exhibit a bi-stability characterized by Up and Down states (fig. 3A-B). Contrary to the system without AHP, in the Up state the dynamics do not exhibit a dominant oscillation band other than *δ* (fig. S2) due to the non imaginary eigenvalues at the Up state attractor. Interestingly, by increasing the network connectivity *J* we can modulate the fraction of time spent in the Up state: for *J* small (*J* = 5.6) the dynamics spends 37% of the time in the Up state, while for *J* = 7.6 it represents 79% (fig. 3C-D and see also fig. S2A-B). Finally, increasing the noise leads to more frequent switches between Up and Down states (fig. S2C-D). To conclude, this model recapitulates the switch between Up and Down states but does not generate a stable *α*-band.

**Figure 3:**
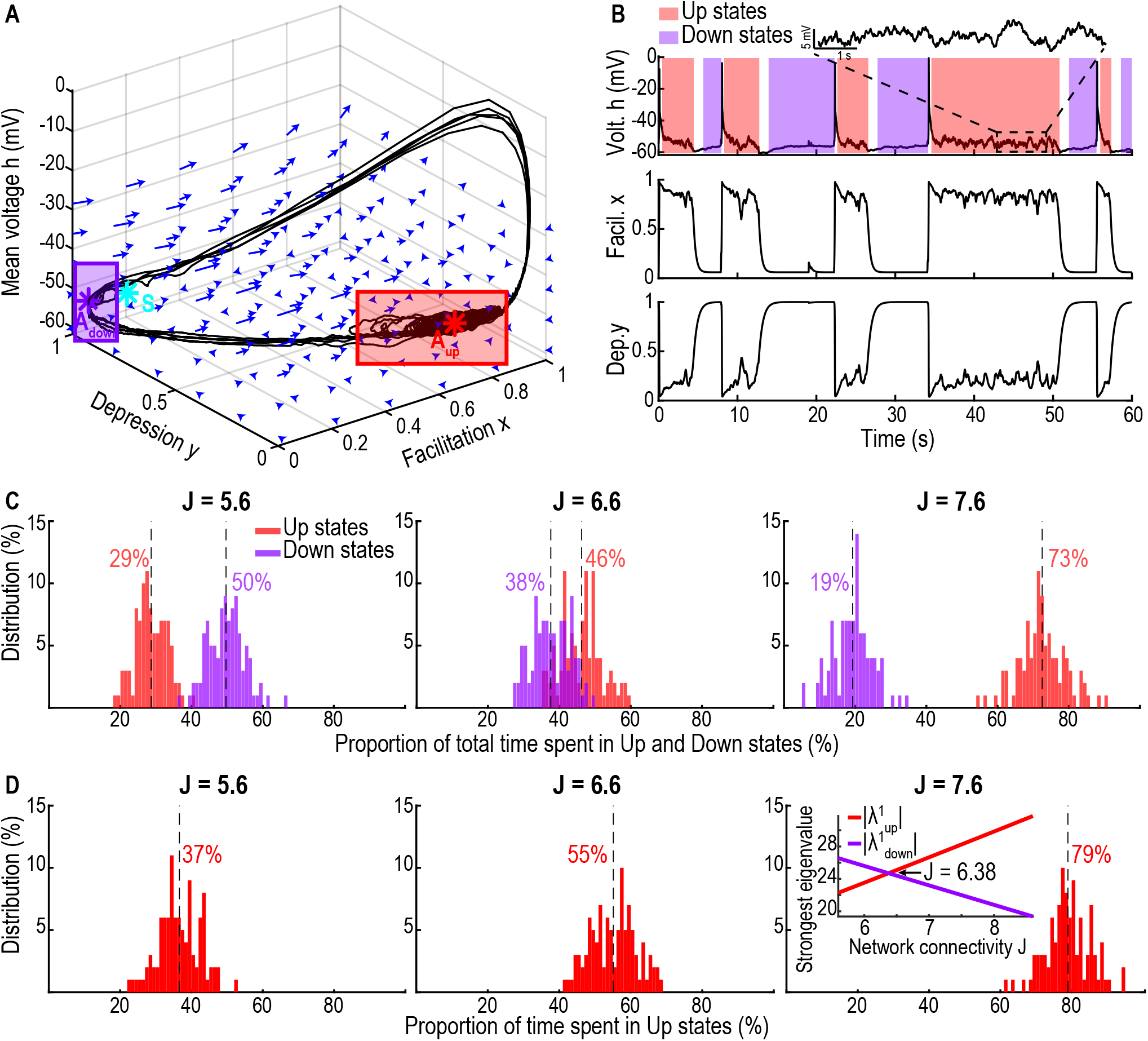
Single population exhibiting Up and Down states. **A.** 3D phase space of system (3) with the two attractors *A_Down_* (purple) and *A_Up_* (red) and the saddle-point *S* (cyan). The purple (resp. red) rectangle shows the subspace of Down (resp. Up) states. **B.** Time series of the mean voltage *h* (upper) showing Down (purple, resp. Up, red) states and an inset during the Up state, facilitation *x* (center) and depression *y* (lower, synaptic plasticity timescales: *τ* = 0.025s, *τ_r_* = 0.5s and *τ_f_* = 0.3s). **C.** Distributions of the fraction of time spent in Down *t_Down_* (purple, resp. Up *t_Up_*, red) states for network connectivity *J* = 5.6 (left), 6.6 (center) and 7.6 (right) for 1000 minutes simulations. Vertical dotted lines indicate mean values. **D.** Distribution of the fraction of the total time spent in Up state 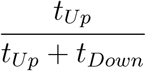 for the three values of *J*. Inset: strongest attractive eigenvalue of the Down state 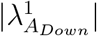 (purple, resp. Up state, 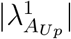, red) with respect to the network connectivity *J*.

#### 1.2.3 Adding a stimulation during the Up states cannot change the oscillation rhythm between Up and Down states in the stable AHP model

Adding an additive input current on the mean voltage *h* during the Up states simulates a situation where the observed network projects an excitatory input on a second network that would send a positive feedback when activated. The second network would only get activated by such stimuli when the first (observed) network is in the Up state. In previous studies with a 2D model (modeling only the firing rate and depression) we showed that such stimulus stabilizes the Up state [24]. Here we ran simulations for the cases with and without AHP where we added a constant input current only when the system was in the Up state (fig. S3). In the case without AHP, the dynamics stays locked in the Up state (fig. S3A), even in the case of a negative feedback current (upper) and the amplitude of the current *I_Up_* does not affect the peak value of the oscillatory band (fig. S3B). In the case with AHP the dynamics is not changed either: the dynamics switches between Up and Down states (fig. S3C) and the proportion of time spent in either Up or Down state is not affected by the value of the current *I_Up_* (fig. S3D).

### 1.3 Modelling the effect of inhibition on the excitatory short term synaptic model with and without AHP

To explore the range of oscillatory behaviors, we connected an inhibitory neuronal network to the excitatory one that could have or not the AHP (see Methods, section 2.3, equations 5). We also added a constant stimulating current *I_i_* on the inhibitory population (fig. 4A).

**Figure 4:**
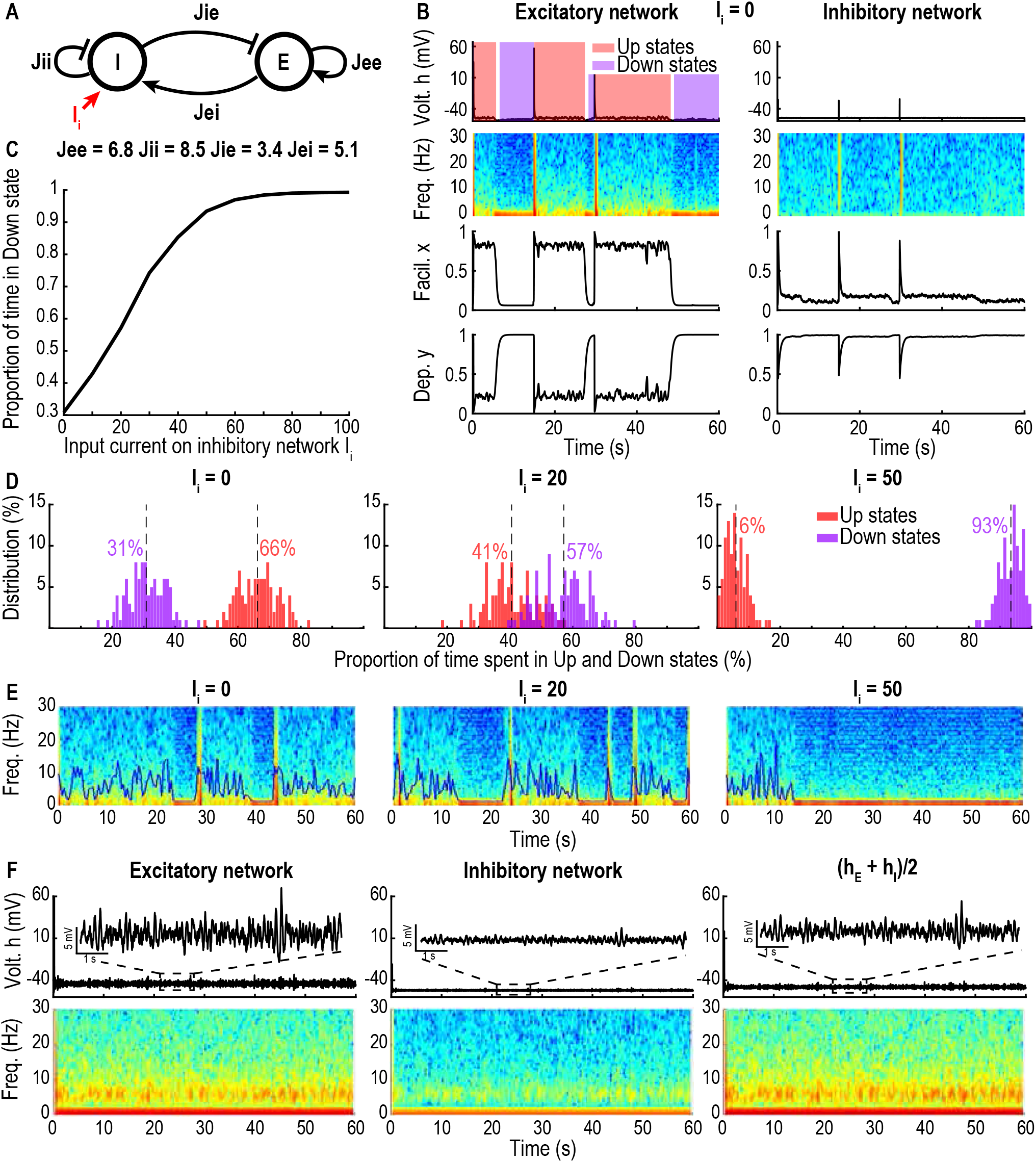
Excitatory-inhibitory network. **A.** Schematic of connectivity between the excitatory (E) and inhibitory (I) networks. **B.** Time series of the excitatory with AHP (left) and inhibitory (right) networks with the spectrograms of *h* for *J_EE_* = 6.8, *J_II_* = 8.5, *J_IE_* = 3.4, *J_EI_* = 5.1, *τ* = 0.01s, *τ_r_* = 0.5s, *τ_f_* = 0.3s and *I_i_* = 0. **C**. Fraction of time spent in the Down state by the excitatory network with respect to *I_i_* ∈ [0,100]. **D**. Distributions of the fraction of time spent in the Down (purple, resp. Up, red) state by the excitatory network for *I_i_* = 0 (left), 20 (center) and 50 (right). E. Spectrograms (60s simulations) of hE for for *I_i_* = 0 (left), 20 (center) and 50 (right) with SEF 95 (blue line). **F**. Time series of the mean voltage *h* (upper) and spectrograms (lower) for the case where the excitatory network does not have AHP and with the timescales *τ* = 0.01s, *τ_r_* = 0.2s, *τ_f_* = 0.12s, showing an *α*-band.

In the case where the excitatory network has AHP the dynamics exhibit switching between Up and Down states (fig. 4B). Interestingly, by increasing the current *I_i_* we modulate the fraction of time spent in Up state by the excitatory system (fig. 4C-D). However, independently of the value of *I_i_* the Up state does not show any persistent *α*-oscillations (fig. 4E). Finally, modulating the current *I_i_* is not sufficient in a dynamics that exhibits the *α*-band to switch between band frequency dominance (fig. 4F).

### 1.4 Two excitatory connected to an inhibitory networks leads to the coexistence of Up and Down states and *α*-oscillations

To define the conditions for which the Up and Down states can coexist with an *α*-band, we explore a model that contains two coupled excitatory components with one inhibitory component (see Methods, section 2.4, equations 6). This investigation is driven by the *α*-oscillation that can be generated by the thalamo-cortical loop (fig. 5A). The thalamo-cortical excitatory subsystem is decomposed into two components *α* and *U/D* connected by reciprocal connections and receives an inhibitory input from the inhibitory subsystem *NR*. The *NR* component sends reciprocal connections to the *U/D* component and can also be activated by an external stimulation *I_i_*. We focus on the sum of the three voltage components because it is the one recorded by EEG. During general anesthesia with propofol, increasing the dose leads to a fragmentation and transient disappearance of the *α*-band. To assess under which conditions this phenomenon could be generated, we followed the same protocol by first investigating the effect of switching off all external stimuli, followed by increasing an injected current to the inhibitory neuronal component to simulate an increase of the propofol concentration.

**Figure 5:**
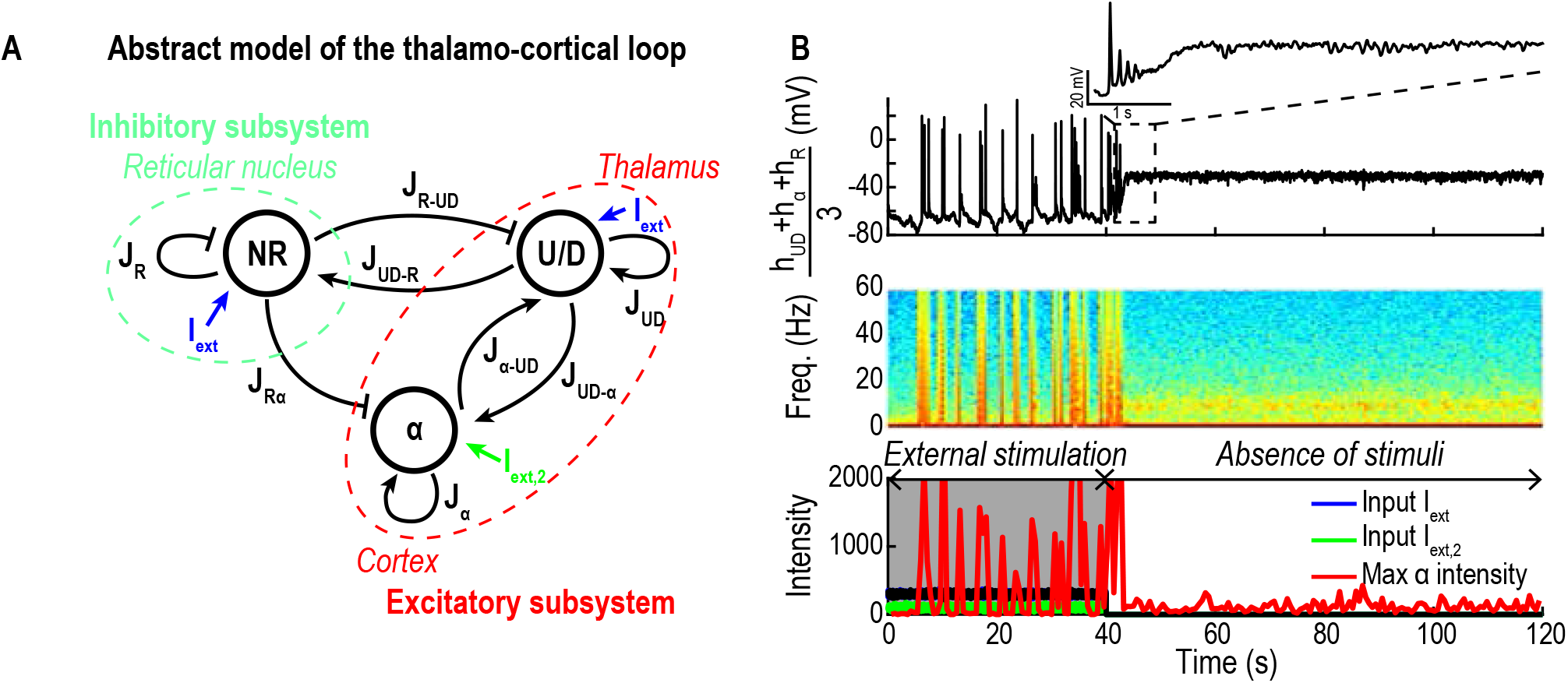
Emergence of the *α*-band following external stimuli suppression. **A.** Connectivity matrix between the two excitatory *α, U/D* and inhibitory (*NR*) network, with the external inputs *I_ext_* (blue) and *I*_*ext*,2_ (green). **B.** Time-series for the sum of the three populations voltage *h_α_* + *h_UD_* + *h_R_* (upper), spectrograms (center) and maximum value of *α* intensity (red) and inputs *I_ext_* (blue and green, lower). The timescale parameters for *U/D* are *τ* = 0.025s, *τ_r_* = 0.5s, *τ_f_* = 0.3s; for *NR* and *α*: *τ* = 0.07s, *τ_r_* = 0.14s, *τ_f_* = 0.086s, *σ_UD_* = *σ_R_* = 6.25 and *σ_α_* = 1.5.

#### 1.4.1 Suppressing external stimuli into two excitatory coupled to an inhibitory network leads to the spontaneous emergence of *α*-oscillations

External stimuli are switch off during the loss of consciousness at the start of a general anesthesia. We modeled here this transition by first adding stimuli modeled as excitatory input current *I_ext_* = 300 + 20ξ (resp. *I*_*ext*,2_ = 100 + 20ξ) where ξ is a Gaussian white noise of mean 0 and variance 1. We applied *I_ext_* and *I*_*ext*,2_ to the three components of the model (fig. 5A, blue and green) for the first 40 seconds of the simulation. To model the beginning of anesthesia, we set the external stimuli *I_ext_* and *I*_*ext*,2_ to zero for the rest of the simulation (fig. 5B). We found that during wakefulness where we set *I_ext_* > 0 from 0 to 40s, there is no dominant oscillatory band in the spectrogram, but after the suppression of the external inputs *I_ext_* = 0 from 40 to 120s, the network stabilizes in the Up state and a dominant *α*-band emerges (fig. 5B). Our simulations suggest that the *α*-band emerges as a stable state once the external stimuli are suppressed, due to the dynamics between the three networks, which reproduces the specific effect of propofol.

#### 1.4.2 Constant low input on inhibition modulates the switching between Up and Down states

We first studied the effect of increasing the inhibitory input current *I_i_* (fig. 6A) on the fraction of time the system spends in Up and Down states. For *I_i_* = 0, we found that the dynamics is characterized by a large proportion of time spent in the Up state (99%) showing persistent *α*-oscillations (fig. 6B). The transition from Up to Down is characterized by a disappearance of the *α*-band, however, the transition from Down to Up is associated with a burst which can either lead to the emergence of an *α*-band or a return to the Down state. By increasing *I_i_* from 0 to 50 and 150 we found that the fraction of time spent in Up states decreases from 99% to 89% and to 4.5% (fig. 6C-E). Each network has a different contribution to the EEG. The *U/D* component (with AHP) shows a fragmented and weak *α*-band while the inhibitory network does not exhibit any particular oscillatory band. Finally, the *α* component (without AHP) exhibits a very strong dominant *α*-band (fig. S4A-B).

**Figure 6:**
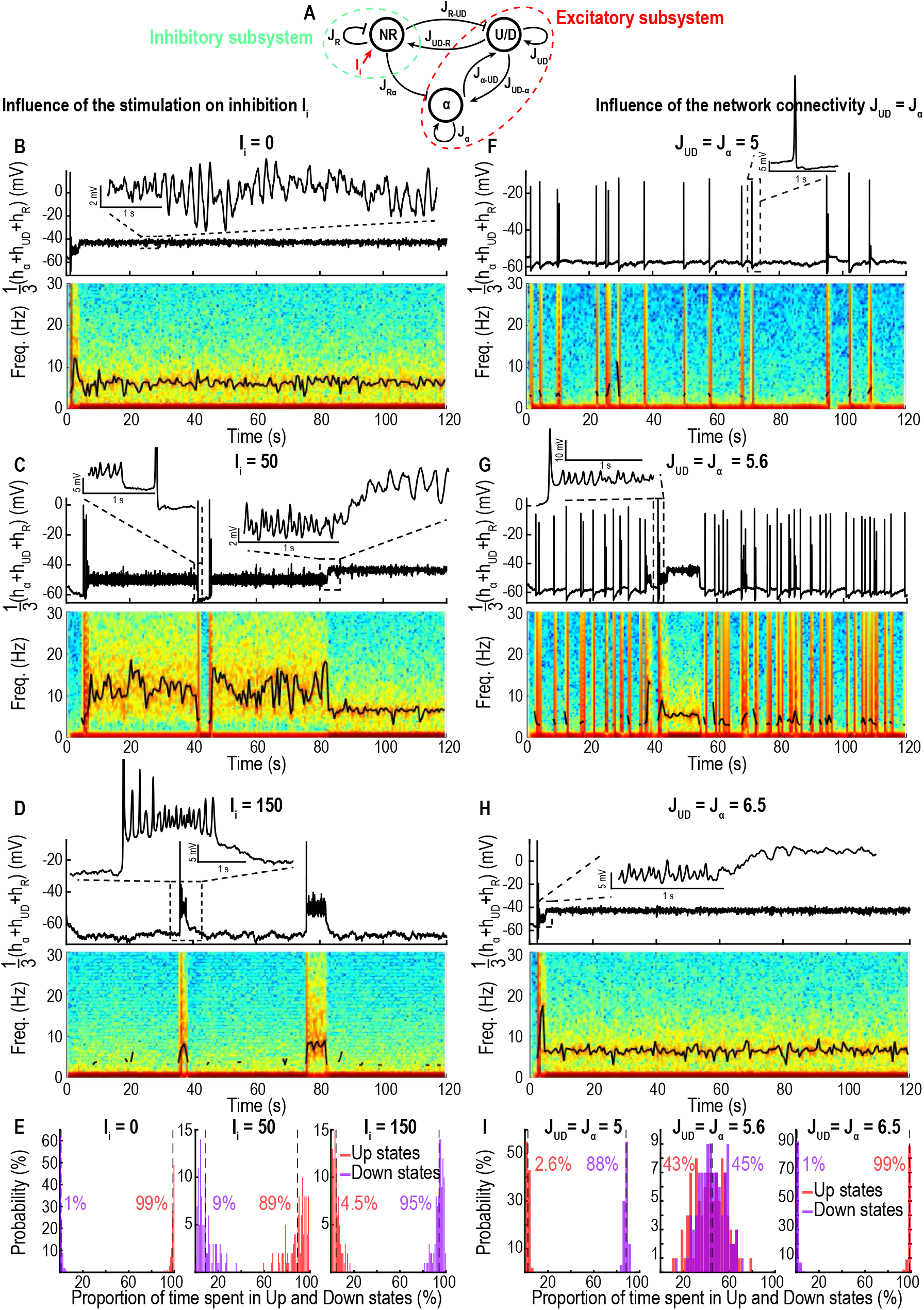
Three compartment model exhibiting Up-Down states and *α*-band in the Up state. **A.** Schematic of connectivity between the two excitatory (*α, U/D*) and inhibitory (*NR*) components. **B-D.** Time-series of the sum of the three populations voltage *h_α_* + *h_UD_* + *h_R_* (upper) and spectrograms (lower) with position of maximum of the oscillatory band (black) for *J_UD_* = *J_α_* = 6.5 and *I_i_* = {0, 50,150}. **E.** Fraction of time spent in the Up and Down states for the three values of *I_i_*. **F-H** Time-series of *h_α_* + *h_UD_* + *h_R_* (Upper) and spectrograms (lower) for *I_i_* = 0 and *J_UD_* = {5,5.6, 6.5}. **I.** Fraction of time spent in the Up and Down states for the three values of *J_UD_* = *J_α_*. Timescales for U/D: *τ* = 0.025s, *τ_r_* = 0.5s, *τ_f_* = 0.3s; for *NR* and *α*: *τ* = 0.005s, *τ_r_* = 0.2s, *τ_f_* = 0.12s

To study the impact of the network connectivity on the emergence of a dominant band, we varied together the intrinsic connectivities *J_UD_* = *J_α_* of both excitatory networks (fig. 6F-H). We found that a small connectivity *J_UD_* = *J_α_* = 5 is associated with a large number of Down states (88%) and transient bursts rarely lead to a stable *α*-band (fig. 6F). By increasing *J_UD_* = *J_α_* to 5.6 and 6.5 the fraction of Up states increases to 43% and 99% respectively (fig. 6I) leading to stable Up states associated with a persistent *α*-band.

#### 1.4.3 Transient responses of the thalamo-cortical model to step and stairs inputs

To study the possible responses of the thalamo-cortical model to propofol bolus and constant increasing we consider a step input (protocol 1) and a stairs (protocol 2) as shown in fig. 7A.

**Figure 7:**
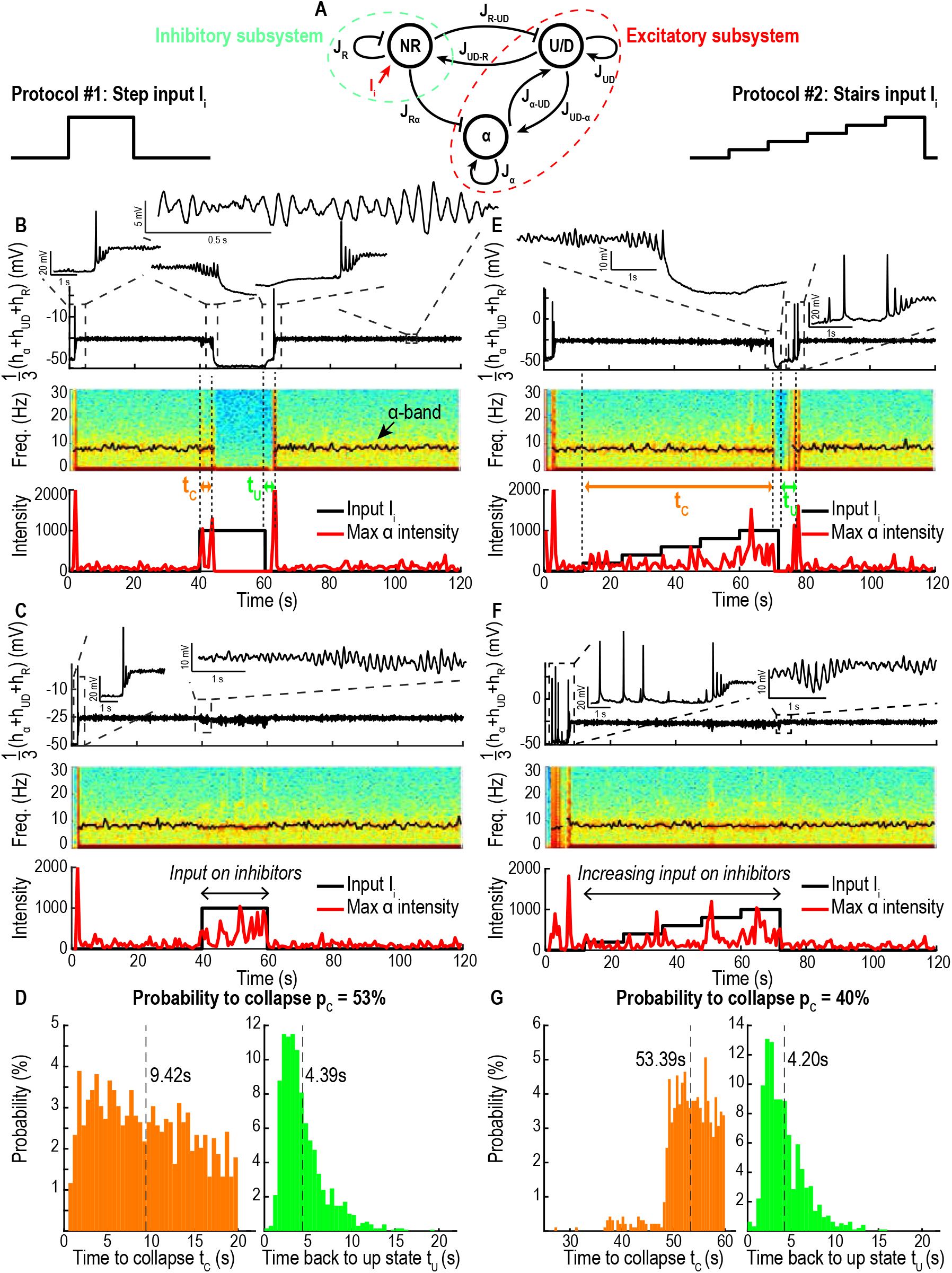
Cases of collapse of the *α*-band. **A.** Schematic of connectivity between the two excitatory (*α* and *U/D*) and inhibitory (*NR*) networks. **B-C**. Time-series of the sum of the three populations voltage *h_α_* + *h_UD_* + *h_R_* (upper), spectrogram (center) with position of maximum of the *α*-band (black, collapsing (B) and persistent (C)), maximum value of *α* intensity (red) and input *I_i_* (black, lower) for a step input. **D.** Distributions of the delay *t_C_* between the beginning of stimulation and the collapse (orange) and delay between the end of the stimulation and the moment the reapparition of the *α*-band (green). **E-G.** Same as B-D for a stairs input. Timescales for *U/D*, *α* and *NR*: *τ* = 0.005s, *τ_r_* = 0.1s, *τ*_f_ = 0.06s

To analyze the response to a step input (protocol 1), we ran simulations for *N* = 2500 iterations lasting *T* = 2min where we simulated a strong injection by a positive input current *I_i_* = 1000 on the inhibitory network (*NR*) lasting *t_i_* = 20s (fig. 7B-C). To quantify the response we collected the statistics of two durations: 1) the duration *t_C_* after which the *α*-band disappears after the step function begins. 2) the duration *t_U_* after which the *α*-band reappears after the end of the step function. Interestingly, for some realizations the *α*-band does not disappear (fig. 7C), we thus characterized this effect by the collapse probability *p_C_*. We found that *p_C_* = 53%, *t_C_* = 9.42 ± 5.36s and *t_U_* = 4.39 ± 2.58s (fig. 7D). The histogram for *t_C_* is characterized by an abrupt decay at 20s confirming that the suppression of the *α*-band can only occur during the stimulation period. However, the time *t_U_* is dominated by an exponential decay, a classical feature of dynamical systems driven by noise over a separatrix.

Each network has a different contribution to the EEG. The *U/D* excitatory network with AHP shows a weak *α*-band while the inhibitory network *NR* only exhibits weak power in the slow *δ*-wave region (≤ 1*Hz*). Finally, the *α* excitatory component without AHP exhibits a very strong dominant *α*-band (fig. S5A-B).

To analyze the effect of a slower increase of the input (stairs function, protocol 2) we ran simulations for *N* = 2500 iterations lasting *T* = 2min where we simulated a stairs increase from *I_i_* = 0 to 1000 on the inhibitory network (*NR*) lasting *t_i_* = 60s (fig. 7E-F). We collected the statistics of the durations *t_C_* (disappearance of the *α*-band) and *t_U_* (reemergence of the *α*-band) as well as the probability to collapse *p_C_*. We found that the probability to collapse is slightly lower in this case *p_C_* = 40%, *t_C_* = 53.39 ± 4.44s and *t_U_* = 4.20 ± 2.43s (fig. 7G). Similarly, the histogram of *t_C_* is characterized by an abrupt decay at 60s confirming that the suppression of the *α*-band can only occur during the stimulation period, while the histogram of *t_U_* is dominated by an exponential decay.

## Discussion

We presented here minimal computational principles based on coarse-grained neuronal network models necessary to generate *α*-oscillations. A single neuronal population driven by synaptic short-term plasticity can illicit oscillations at a defined frequency, which directly depends on the value of the network connectivity: a higher connectivity generates faster oscillations (fig. 2B-C). Interestingly, we show here that the *α*-oscillations results from the combination of network connectivity, synaptic and biophysical properties, leading to a focus attractor, around which the stochastic mean population voltage oscillates in the phase-space (fig. S7A-B). Moreover, we showed that spontaneous switching between Up and Down states in a single neuronal population is modulated by AHP and also that the network connectivity controls the proportion Up vs Down states: a higher connectivity *J* results in a dominant percentage of time spent in Up states (fig. 3C-D). The stability of the oscillations during Up states for a population without AHP could result from the intrinsic network regulation: indeed, interactions between hundreds of inhibitory interneurons and hippocampal pyramidal excitatory neurons can redistribute the firing load to maintain the oscillation frequency even when up to 25% of the synapses are deactivated [32].

When we added excitatory input current on the inhibitory coupled to excitatory population, the proportion of time spent in Up states decreases and, after reaching a threshold value (*I_i_* = 60), the network becomes completely silenced, characterized by Down states only (fig. 4C-D). When coupling two excitatory and one inhibitory neuronal population (fig. 5A), *α*-oscillations, generated by the excitatory component without AHP, co-existed with spontaneous switching between Up and Down states induced by the excitatory population with AHP, as summarized in fig. 8. Stimulating the inhibitory population induces the fragmentation of the *α*-band by modulating directly the proportion of Up vs Down states (fig. 6B-E).

**Figure 8:**
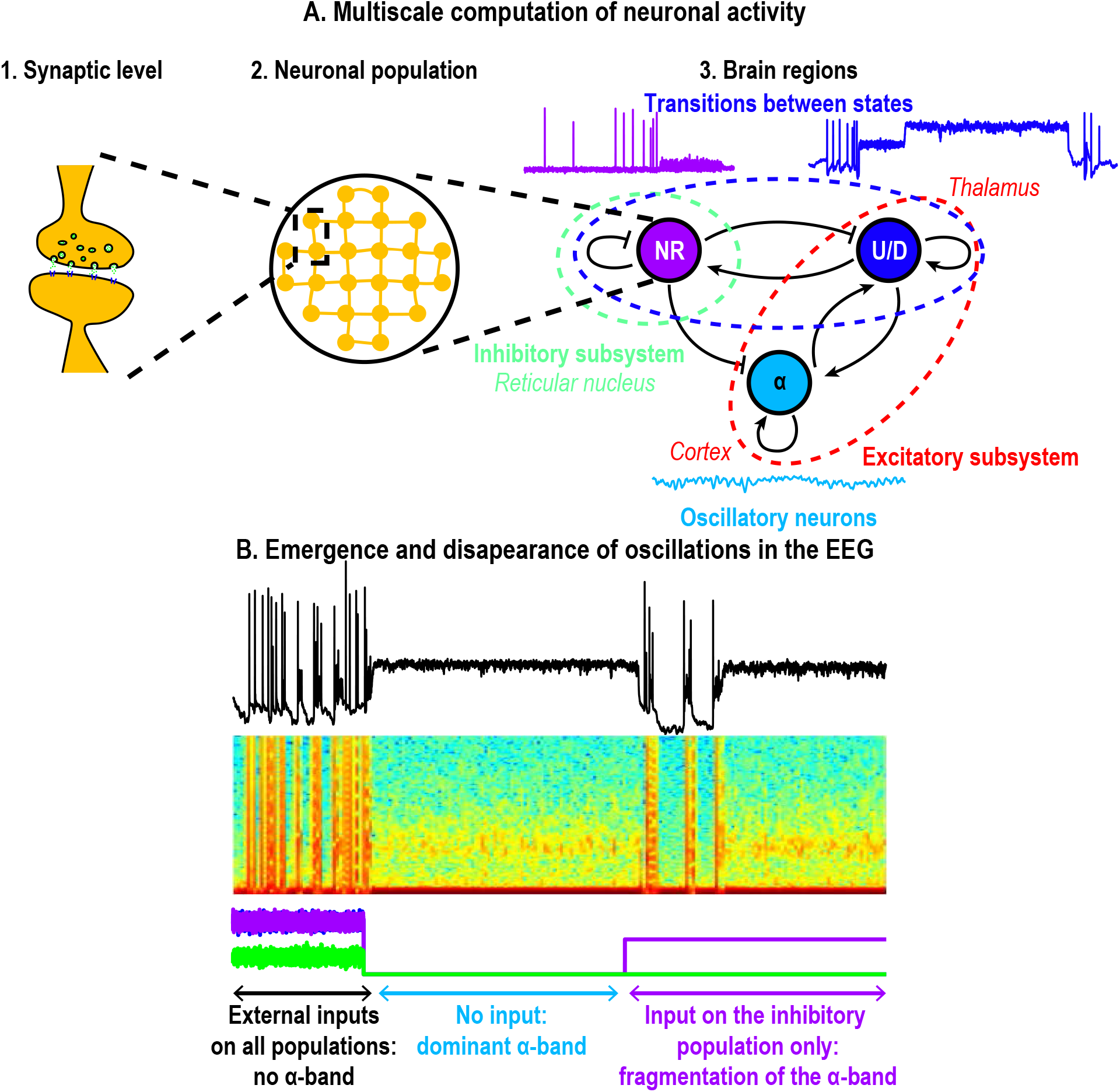
Computational principle underlying the *α*-band dynamics. **A.** Multiscale models 1. synaptic level: short-term facilitation and depression mechanisms, 2. neuronal population: mean voltage fluctuations and 3. interactions between different brain regions (thalamus and cortex, models 5 and 6) resulting in the different activity patterns observed in EEG at a global scale. **B.** *α*-band emerges in the Up-state due to noise but can be altered by external inputs.

Finally, we suggest that synaptic noise has two main roles on the network properties: 1) increasing the noise intensity stabilizes the *α*-band (fig. 2D) and 2) when an external stimulation is applied to the inhibitory system in a step or stairs input, the network can react with opposite behavior: either the network activity collapses, leading to a suppression of all oscillatory bands in the EEG or a stable persistent *α*-band emerges during the entire stimulation (fig. 7). Finally, we propose that three connected neuronal populations are sufficient to generate an *α*-oscillation that could be fragmented by increasing the inhibitory pathway, as suggested during general anesthesia [7]. The present model could be generalized to study the emergence and disappearance of other oscillations such as the *θ*-oscillations occurring during REM sleep [2,33,34].

### Modeling the dynamics of the *α*-band

The origin of the *α*-band [14] remains unclear. Early modeling efforts using the Hodgkin-Huxley framework [30] suggested a key role of ionic currents such as sodium, potassium currents, low threshold calcium, AHP and synaptic currents (GABAs, AMPA) that could reproduce various patterns of oscillations [19] as well as initiation, propagation and termination of oscillations (see also [35]). By varying the GABA conductances similar models could reproduce the dominant *α*-oscillation observed in propofol anesthesia [20,21]. Indeed the GABA conductance regulates the firing frequencies and the synchronization of pyramidal neurons [36]. In contrast with models based on ionic conductances, in the present model, based on short-term synaptic plasticity driven by noise, the *α*-band is generated only when the mean voltage is in the Up state, suggesting that the ionic mechanisms are not necessary to generate the *α*-band, but contributes to the termination of the Up states. Furthermore, adding an input current to the inhibitory population allows to generate transitions between Up and Down states (fig. 4C-D and 6B-E). In addition, we found that the *α*-band can be stabilized by increasing the noise amplitude, while the peak frequency of the *α*-band was unchanged (fig. 2D-E). Thus, we propose that the synaptic noise could be responsible for the stabilization of the *α*-band, as quantified by our fragmentation analysis (fig. 2D-E). Finally, we reported a second mechanism responsible for an *α*-band fragmentation associated with the transition between Up and Down states (fig. 6).

Interestingly, the *α*-band is persistent in young subjects and becomes sparser with age [37]. Possibly, a higher neuronal activity (in younger subjects) leads to higher extracellular potassium which, in turn, increases the synaptic noise [38]. Another possible mechanism for fragmenting the *α*-band could involve the metabolic pathway, when the ATP coupled to the sodium concentration is decreased: during a burst, a high sodium concentration depletes ATP that deregulates the potassium current and thus leads to a phase of iso-electric suppression [10].

### Relation between Up and Down states and the *α*-band

Neuronal networks exhibit collective transitions from Up to Down states [15, 16, 22]. We reported here that the *α*-oscillations is only generated when the neuronal ensemble is in the Up state. Interestingly, we could not generate, in a single neuronal population, at the same time this *α*-oscillation and the Up-Down states transitions. Rather we needed a minimum of two coupled excitatory neuronal populations. We reported here that the fraction of time spent in Up and Down states depends on the level of synaptic connectivity (fig. 3C-D and 6F-I). However, by adding an inhibitory network, we were able to modulate the proportion of time spent in Up vs Down states by changing the input stimulation current (fig. 4C-D and 6B-E) on this inhibitory population without varying the connectivity. The mechanism is feasible because the inhibitory input on both excitatory populations allows to destabilize the Up state and thus increase the transitions to Down state modulating the overall fraction of time spent in the Up states. To conclude, in the extreme case where the Down states are dominant, the overall voltage dynamics resemble iso-electric suppressions without the need to account for a metabolic stress [39, 40].

### Predictions and limitations of the model to interpret the *α*-band during general anesthesia

The physiological mechanisms leading to the emergence of the *α*-band shortly after propofol injection during general anesthesia remains unclear [7,41]. Possibly, during wakefulness, the amount of external stimuli suppresses the emergence of *α*-oscillations [2]. When the external stimuli ceases with propofol injection, the *α*-oscillation could become dominant (fig. 5B). The present model suggests that the initial state represents an already anesthetized brain where the neuronal networks do not receive any external stimuli, leading to the spontaneous emergence of the *α*-band.

General anesthesia needs to be sustained over the whole course of a surgery and thus controlling optimally the anesthetic injection to prevent cortical awareness or a too deep anesthesia remains a difficult problem [37]. Population models such as the one presented here could be used to test different activation pathways of anesthetic drugs. The present model accounts for the appearance an iso-electric suppression [42] in the EEG of the order of a few seconds (fig. 7B,E) induced by increasing transiently the hypnotic. We predicted here that the fragmentation of the *α*-band results from a shift between Up and Down states dominance that could be tested with in vivo experiments. It would be interesting to further account for longer term consequences of an anesthetic input (several minutes). Indeed, the causality between *α*-suppressions and burst-suppressions [11] remains unexplained, suggesting that this relation could involve other mechanisms than the ones we modeled here based on synaptic plasticity, AHP and network connectivity.

## 2 Methods

### 2.1 Fragmentation level of an oscillatory band

The emergence of the *α*-rhythm is characterized by a continuous band in the range [8–12]Hz in the spectrogram of the EEG. We define here a measure for the persistence in time of this band. Once we detected the peak spectral value *S_α_*(*t*) of the spectrogram as the highest power value in the extended range *α_min_* = 4 – *α_max_* = 16 Hz when the condition *S_α_*(*t*) > *T_α_*, then we consider that the band is present and attribute a value *x_pr_*(*t*) = 1, otherwise *x_pr_*(*t*) = 0. When the time interval between 0 and *T* is divided into N bins at times *t_k_*, the presence of the *α* band is defined by

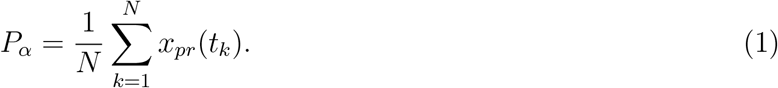

The persistence level *P_α_* measures the proportion of time where the *α*-band is present.

To further quantify the fragmentation level, we introduce the disruption number *D_α_* that counts the number of time per minute the peak spectral value *S_α_*(*t*) goes under the threshold *T_α_*.

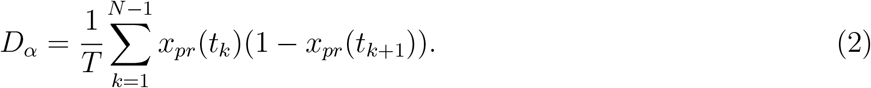

We call the fragmentation level the pair *F_α_* = (*P_α_, D_α_*) (fig S6). For the human EEG data (fig. 1), we used a bin size *w* = 0.5*s* and a threshold value *T_α_* = 1.5*dB*. For the simulated data, we use the same bin size and *T_α_* = 10.

### 2.2 Modeling a single neuronal population based on synaptic depression-facilitation dynamics

For a sufficiently well connected ensemble of neurons, we use a mean-field system of equations to study bursting dynamics, AHP and the emergence of Up and Down states. This stochastic dynamical system consists of three equations [23, 26, 29] for the mean-field variable *h*, the depression *y*, and the synaptic facilitation *x*:

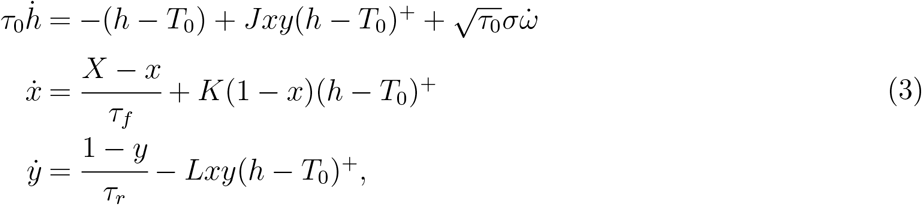

where *h*^+^ = *max*(*h*, 0) is the population mean firing rate [24]. The term *Jxy* reflects the combined synaptic short-term dynamics with the network activity. The second equation describes facilitation, and the third one depression. The parameter *J* accounts for the mean number of synaptic connections per neuron [23,43]. We previously distinguished [26] the parameters *K* and *L* which describe how the firing rate is transformed into synaptic events that are changing the duration and probability of vesicular release respectively. The time scales *τ_f_* and *τ_r_* define the recovery of a synapse from the network activity. We account for AHP with two features: 1) a new equilibrium state representing hyperpolarization after the peak response of the burst 2) two timescales for the medium and slow recovery to the resting membrane potential to describe the slow transient to the steady state. Finally, 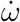 is an additive Gaussian noise and *σ* its amplitude, representing fluctuations in the mean voltage.

In the case of a neuronal network that does not exhibit AHP the resting membrane potential is constant *T*_0_ = 0 and *τ*_0_ = *cst* ∈ [0.005, 0.025]s. However, for a population showing AHP after the bursts, the resting membrane potential *T*_0_ and the recovery time constant *τ*_0_ of the voltage *h* are defined piece-wise as follows:

- *τ*_0_ = *τ* and *T*_0_ = 0 in the subspace Ω*_fast_* = {*y* > *Y_AHP_* and *h* ≥ *H_AHP_*}, which represents the fluctuations around the resting membrane potential during the down state and the burst dynamics.
- *τ*_0_ = *τ_mAHP_* and *T*_0_ = *T_AHP_* < 0 in the subspace 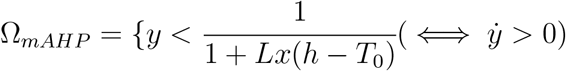 and *y* < *Y_h_*}. This part of the phase-space defines the moment when the hyperpolarizing currents at the end of the burst become dominant and force the voltage to hyperpolarize.
- *τ*_0_ = *τ_sAHP_* and *T*_0_ = 0 in the subspace 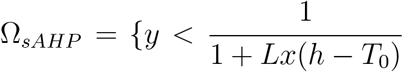 and (*Y_AHP_* < *y* or *h* < *H_AHP_*)}, which represents the slow recovery to resting membrane potential.

The threshold parameters defining the three phases are *Y_h_* = 0.5, *Y_AHP_* = 0.85 and *H_AHP_* = –7.5. In this study, we varied the network connectivity parameter *J* ∈ [5.6, 8.6] and all other parameters are described in Table 1, SI.

To convert the mean-field variable *h* into a mean voltage 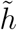 in mV, we use the following conversion

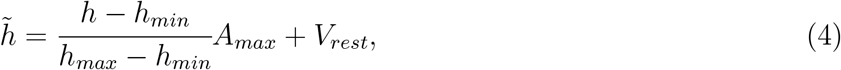

where *V_rest_* = –70 mV. We identified *h_min_* = –100 and *h_max_* = 1200 based on numerical simulations and chose *A_max_* = 200*mV* according to the classical amplitude of intracellular recordings.

### 2.3 Two-populations model of the thalamo-cortical loop

To model the interactions between one excitatory *E* and inhibitory *I* neuronal network, we coupled two systems of equations (3)) as follows:

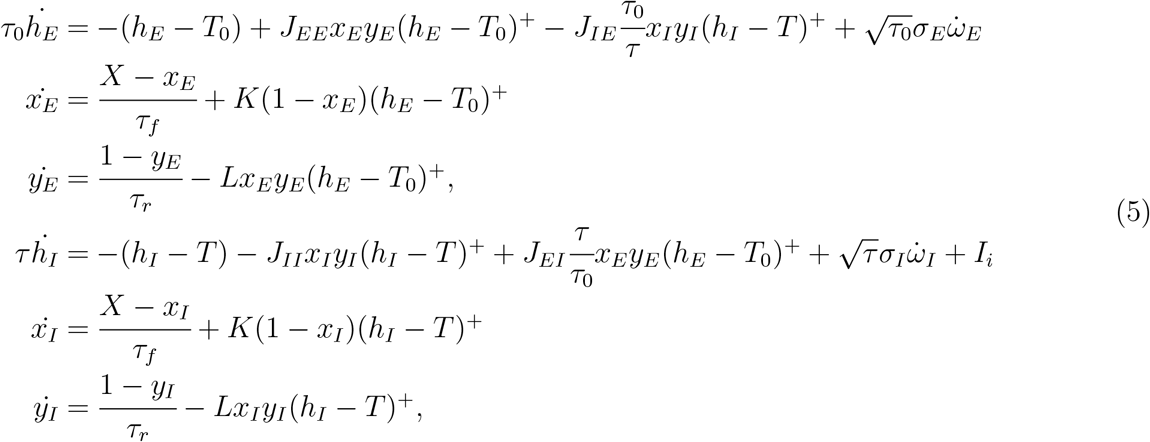

where *τ*_0_ and *T*_0_ for the excitatory population can either be constant, in the absence of AHP or defined piece-wise when it is present, as already discussed in subsection 2.2. The inhibitory population is always modeled without AHP and thus *τ* is constant and *T* = 0. All other parameters are described in the central columns called “2 populations” of Table 1, SI.

### 2.4 Three connected neuronal populations to model the thalamo-cortical loop

To model the thalamo-cortical loop, we connected three neuronal networks. One excitatory network driven by AHP generates the Up-Down state dynamics (referred to as *U/D* in figs. 5, 6 and 7). The second excitatory network is not driven by AHP and is referred as *α* in figs. 5, 6 and 7. Both networks are coupled with an inhibitory one (called *NR*), which does not exhibit any AHP. The equations extend the case of two neuronal networks presented in subsection 2.3 and the connectivity matrix with 9 elements is presented in Table 1, SI. The overall system of equation is

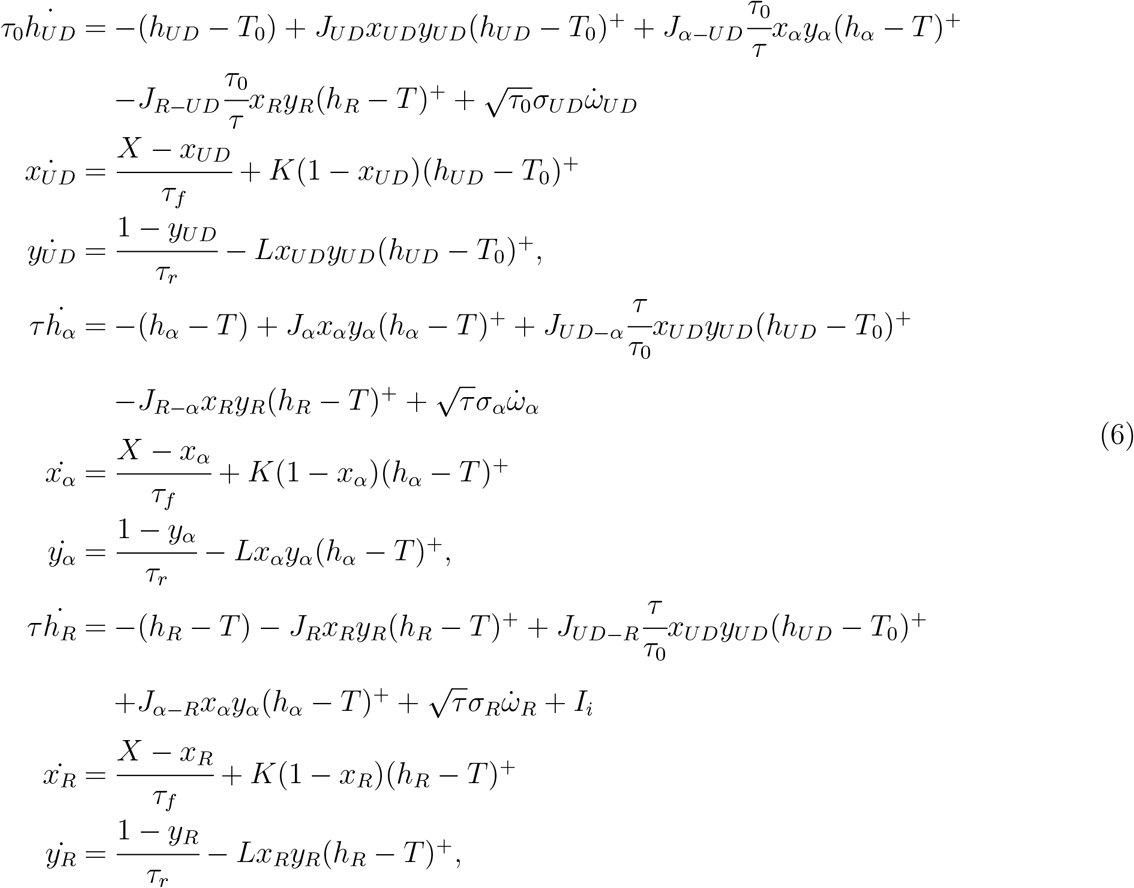

where *τ*_0_ and *T*_0_ for the first excitatory population *U/D* are defined piece-wise in part 2.2 and all other parameters are given in Table 1, SI (right columns: “3 populations”).

### 2.5 Origin of oscillations in the Up state

We study here the origin of the oscillations observed in the spectrograms of *h* in relation with the Up and Down states.

#### 2.5.1 Oscillations around the Up state attractor for a neuronal population without AHP

In the absence of AHP, the focus attractor *A_Up_* has two complex conjugated eigenvalues. Thus the deterministic dynamics oscillates around the point *A_Up_* at a frequency

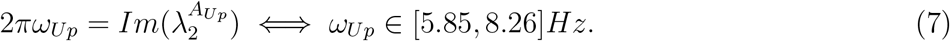

which corresponds to the dominant spectral band observed in fig. 2. The oscillation eigenfrequency *ω_Up_* further depends on the network connectivity *J* (fig. 2A-B), but not on the noise amplitude (fig. 2C-D). Note that the noise allows to generate persistent oscillation compared to the case of the pure deterministic system. Finally, increasing the noise amplitude stabilizes the *α*-band (fig. 2C-D).

Note that if we take *τ* = 0.025*s*, *τ_r_* = 0.5*s* and *τ_f_* = 0.3*s*, then the eigenvalues of *A_Up_* become 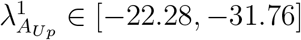 and the complex-conjugate eigenvalues

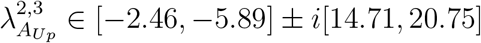

leading to an eigenfrequency *ω_Up_* ∈ [2.34, 3.30]Hz (fig. S1) which explains the disappearance of the dominant *α*-band in this case (see also fig. S7C).

Finally, since 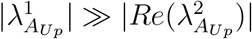 the dynamics is very anisotropic and the oscillations are confined in a 2D manifold (fig. S7A.1-A.3, light red trajectories).

#### 2.5.2 The Up state stability is due to multiple re-entries in its basin of attraction

To explain the locking in the Up state, we recall that the stochastic trajectories starting inside the basin of attraction of the Up state can cross the separatrix Γ and fall into the Down state. However, because the deterministic vector field of system (3) is very shallow near Γ, the additive noise on the *h* variable can push the trajectories back into the Up state, where the field is stronger, and thus the trajectory is brought in a neighborhood of *A_Up_* and continues oscillating, as shown in fig. S7B (see inset).

To explain the other frequencies (than the eigenfrequency *ω_Up_*) observed in the spectrum of *h* (fig. S7C), we note that when a trajectory falls back in the Up state, it can produce a longer or shorter loop depending on its initial distance to the attractor *A_Up_*. These oscillations between the two basins of attraction define stochastic oscillations that contribute to the spectrogram of *h*.

#### 2.5.3 Oscillations between Up and Down state in a neuronal population containing an AHP component

For a neuronal network with an AHP component, the Up state has only real negative eigenvalues (fig. S8C), thus no oscillations are expected near the attractor. However, the presence of a slow AHP component (fig. S8A-B pink) can push the dynamics into the Down state, as opposed to the case without AHP. Finally, in the Down state, the trajectories fluctuate with the noise until they escape. Once trajectories cross the separatrix Γ, they follow an almost deterministic path close to that of the unstable manifold of S fig. S8A-B grey) showing a long excursion in the phase-space before falling back near the attractor *A_Up_*. This dynamics explains the recurrent switches between Up and Down states.

