## Supplementary material for "Emergence and fragmentation of the alpha-band driven by neuronal network dynamics": SI: methods, figures

### Up and Down states occurring in neuronal networks regulate the emergence and fragmentation of the alpha-band

#### 1 Supplementary results

Table 1 summarizes the parameters used for all the simulation results presented in the main text and in the following supplementary figures.

| | 1 population | | 2 populations | | 3 populations | | |
|---|---|---|---|---|---|---|---|
| | no AHP | AHP | no AHP & I | AHP | same E | no AHP & I | AHP |
| $\tau$ | 0.005 ($\alpha$) - 0.01s ($\theta$) | 0.025s | 0.005 ($\alpha$) - 0.01s | 0.025s | 0.005s | 0.005-0.07s | 0.025s |
| $\tau_r$ | 0.2 - 0.5s | 0.5s | 0.2 - 0.5s | 0.5s | 0.1s | 0.1 -0.2s | 0.5s |
| $\tau_f$ | 0.12 - 0.3s | 0.3s | 0.12 - 0.3s | 0.3s | 0.06s | 0.06 - 0.12s | 0.3s |
| $\tau_{mAHP}$ | | 0.3s | | 0.12s | 0.06s | | 0.12s |
| $\tau_{sAHP}$ | | 1s - 10.5s | | 1s | 0.5s | | 1s |
| $J_{E_1E_1}$ | 5.6 - 8.6 | | 6.8 | | 5.6 | 6.5 | |
| $J_{E_1I}$ | | | 5.1 | | 5.6 | 6.5 | |
| $J_{IE_1}$ | | | 3.4 | | 4.48 | 16.25 | |
| $J_{II}$ | | | 8.5 | | 5.6 | 3.25 | |
| $J_{E_1E_2}$ | | | | | 2.8 | 1.3 | |
| $J_{E_2E_1}$ | | | | | 1.12 | 1.3 | |
| $J_{E_2I}$ | | | | | 0 | 0 | |
| $J_{IE_2}$ | | | | | 4.48 | 16.25 | |
| $J_{E_2E_2}$ | | | | | 4.2 | 6.5 | |
| $\sigma$ | 5 - 15 | | 2.75 ($\sigma_I$) 5.5 ($\sigma_E$) | | 10 ($\sigma_T$) 3 ($\sigma_{C,R}$) | 2.5 ($\sigma_{T,C,R}$) | |
| $T_{AHP}$ | | -30 | | -30 | -30 | | -30 |
| $K$ | 0.5 Hz | | | | | | |
| $L$ | 0.3 Hz | | | | | | |
| $X$ | 0.06 | | | | | | |

Table 1: Models 1 (1 population), 2 (2 populations) and 3 (3 populations) parameters (see Main text, Methods). For models (2) and (3), the inhibitory population is always without AHP and excitatory populations can be with or without AHP. For model (3) $E_1$ corresponds to the network with AHP ($U/D$), and $E_2$ to the network without AHP ($\alpha$).

[1]Sorbonne University, Pierre et Marie Curie Campus, 5 place Jussieu 75005 Paris, France.
[2]Group of Applied Mathematics and Computational Biology, École Normale Supérieure, France.

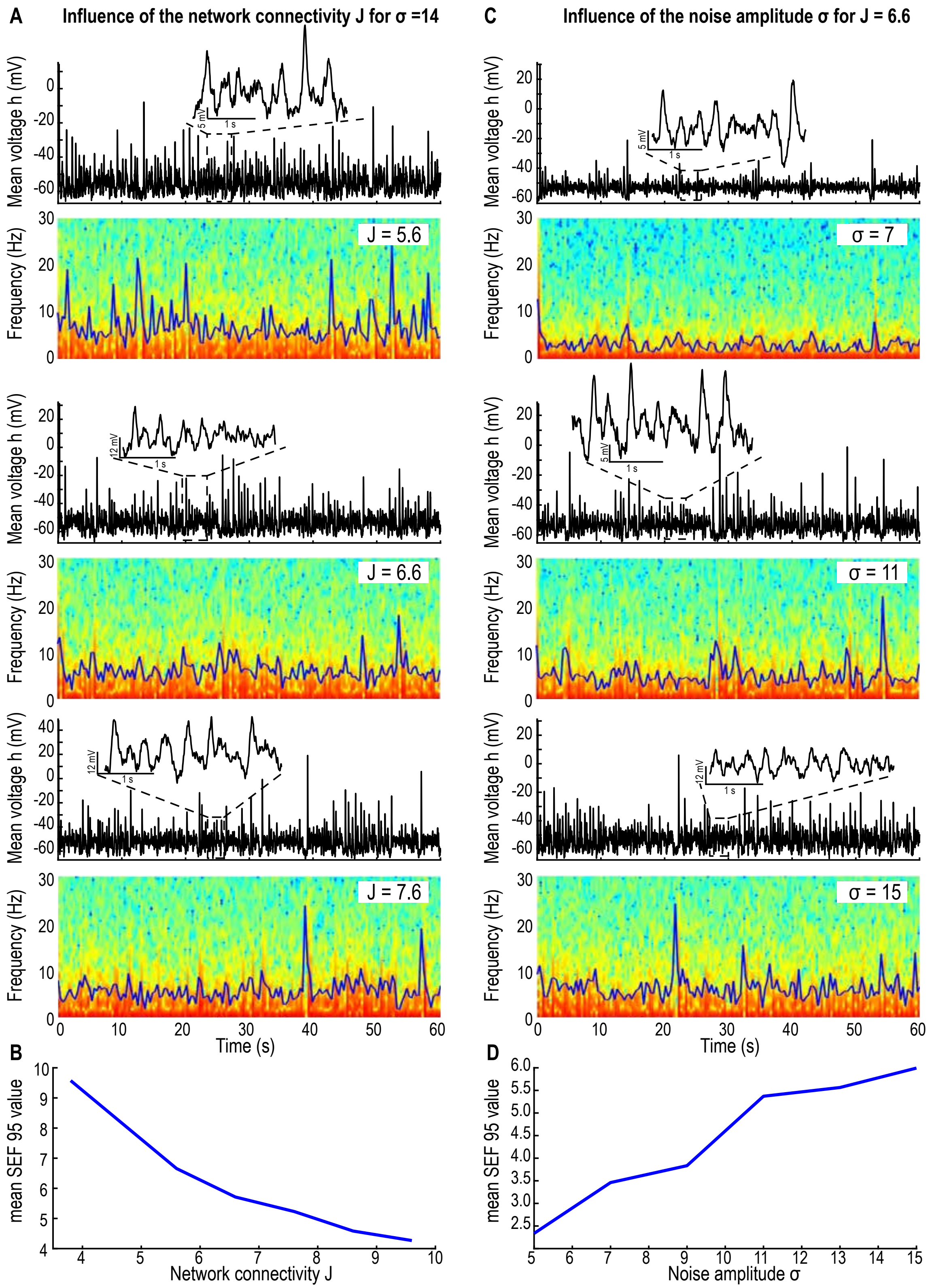


Figure S1: **Effect of network connectivity $J$ and noise amplitude $\sigma$ on model (1) without AHP. A.** Time-series and spectrograms of $h$ (60s simulations) with SEF95 (blue curve) for $J = 5.6$ (upper), 6.6 (center) and 7.6 (lower). **B.** Mean value of the SEF95 for $J \in [3.8, 10]$. **C.** Time-series and spectrograms of $h$ (60s simulations) with SEF95 (blue curve) for $\sigma = 7$ (upper), 11 (center) and 15 (lower). **D.** Mean value of the SEF95 for $\sigma \in [5, 15]$. Synaptic plasticity timescales: $\tau = 0.025$s,$\tau_r = 0.5$s and $\tau_f = 0.3$s.

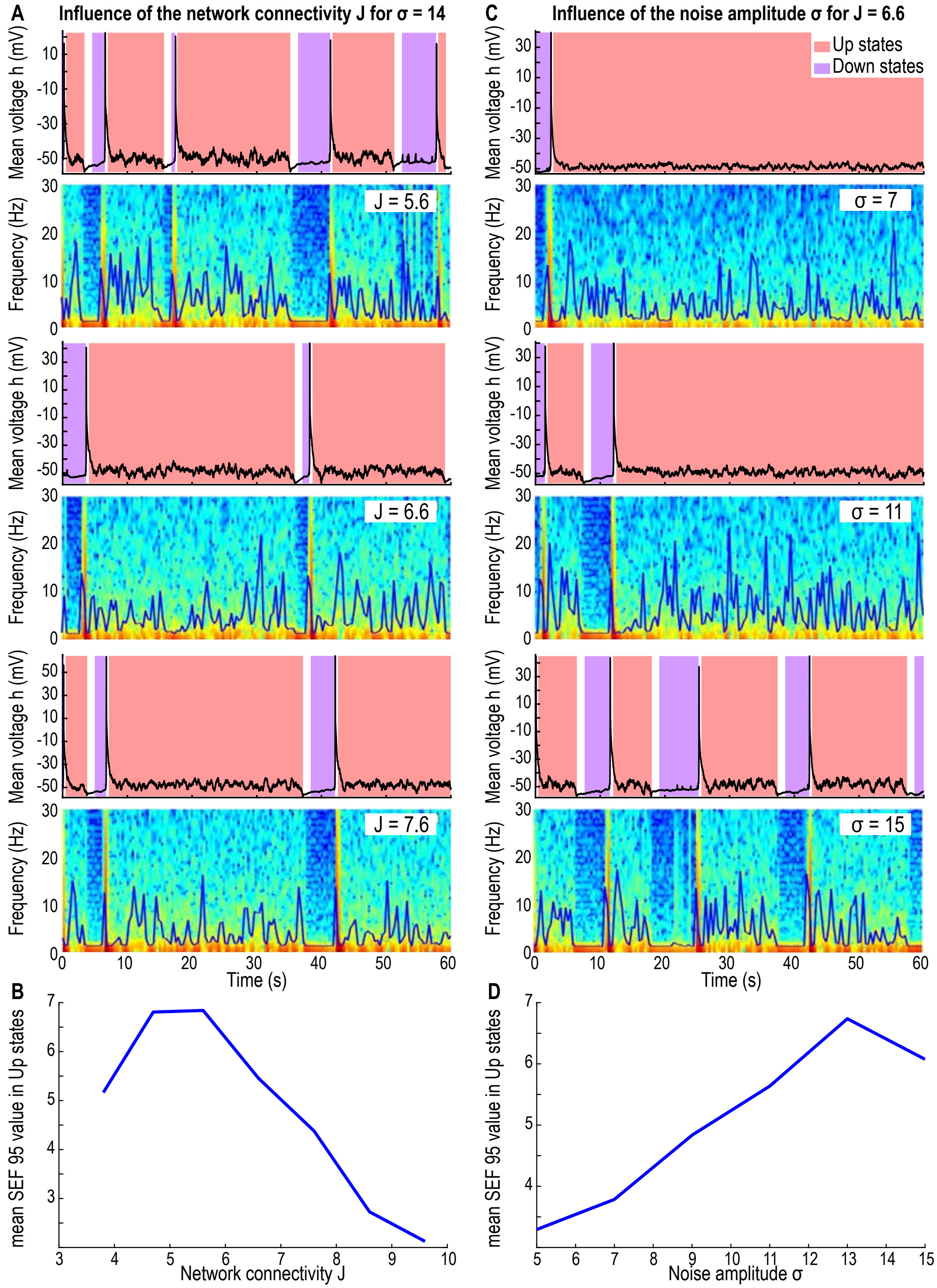


Figure S2: **Effect of network connectivity $J$ and noise amplitude $\sigma$ on model (1) with AHP. A.** Time-series and spectrograms of $h$ (60s simulations) with SEF95 (blue curve) for $J = 5.6$ (upper), 6.6 (center) and 7.6 (lower). **B.** Mean value of the SEF95 in the upstates for $J \in [3.8, 10]$. **C.** Time-series and spectrograms of $h$ (60s simulations) with SEF95 (blue curve) for $\sigma = 7$ (upper), 11 (center) and 15 (lower). **D.** Mean value of the SEF95 in the upstates for $\sigma \in [5, 15]$. Synaptic plasticity timescales: $\tau = 0.025$s,$\tau_r = 0.5$s and $\tau_f = 0.3$s.

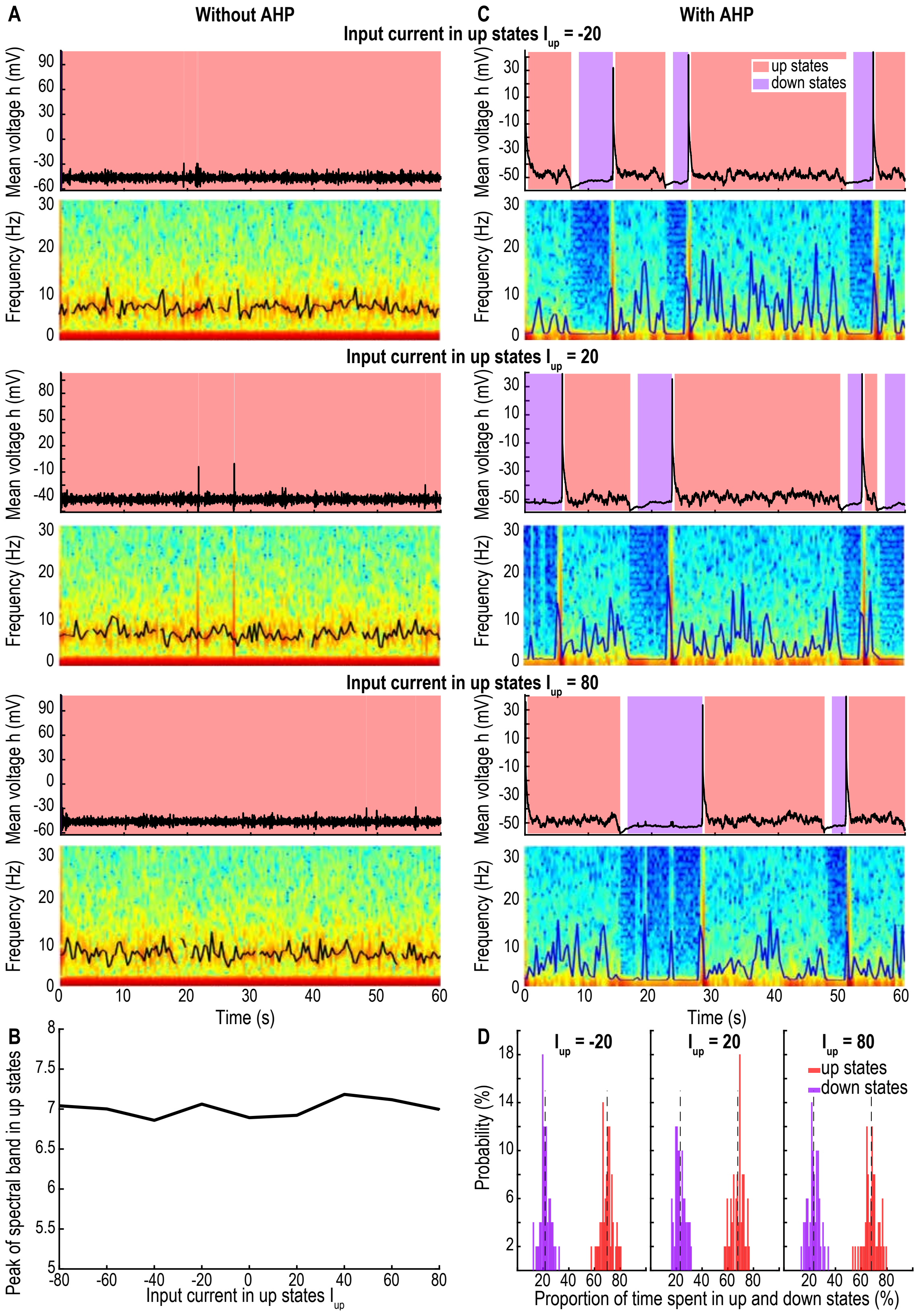


Figure S3: **Effect of an input current $I_{up}$ during the up states in model (1). A.** Time-series and spectrograms of $h$ (60s simulations, model (1) without AHP with $J = 6.6$, $\sigma = 10$, $\tau = 0.01$s,$\tau_r = 0.2$s and $\tau_f = 0.12$s), with peak value of the oscillatory band, (black curve) for $I_{up} = -20$ (upper) 20 (center) and 80 (lower). **B.** Mean peak value of the oscillatory band for $I_{up} \in [-80, 80]$. **C.** Time-series and spectrograms of $h$ (60s simulations, model (1) with AHP with $J = 6.6$, $\sigma = 14$, $\tau = 0.025$s,$\tau_r = 0.5$s and $\tau_f = 0.3s$). **D.** Proportion of time spent in up vs down states for $I_{up} = \{-20, 20, 80\}$ ($N = 50$ simulations of $T = 5$min, model (1) with AHP).

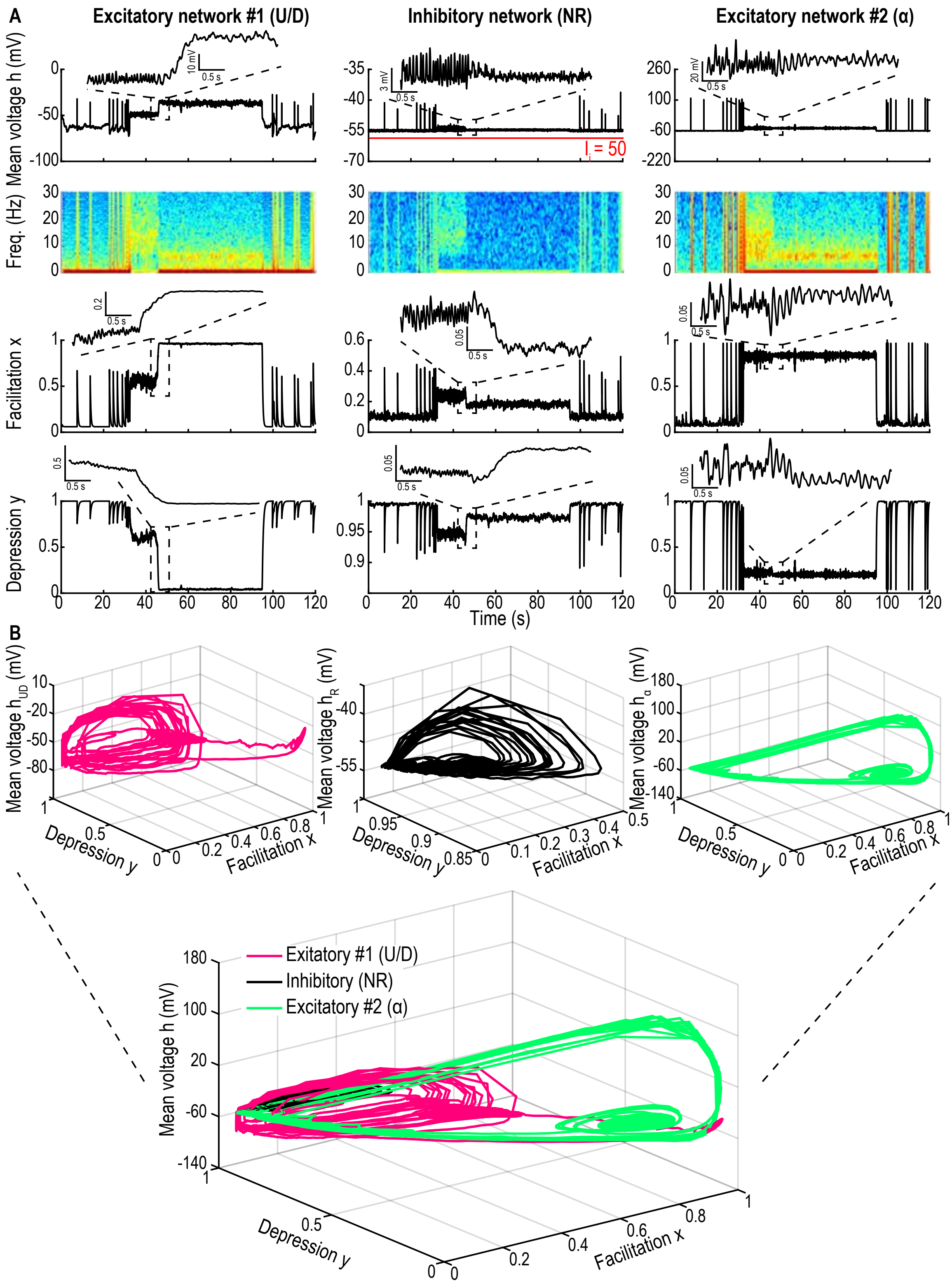


Figure S4: **Contribution of the three components of model (3) for a constant input. A.** Time-series of mean voltage $h$, spectrogram, facilitation $x$ and depression $y$ of system 3 (120s simulations) for the excitatory network with AHP ($U/D$, left: $\tau = 0.025$s, $\tau_f = 0.3$s,$\tau_r = 0.5$s), the inhibitory network ($NR$, center) and the excitatory network without AHP ($\alpha$, right: $\tau = 0.005$s, $\tau_f = 0.12$s, $\tau_r = 0.2$s) with a constant input $I_i = 50$ on the inhibitory network (red line). **B.** Trajectories in the $h - x - y$ phase space of each component ($U/D$, pink, left, NR black, center and $\alpha$, green, right).

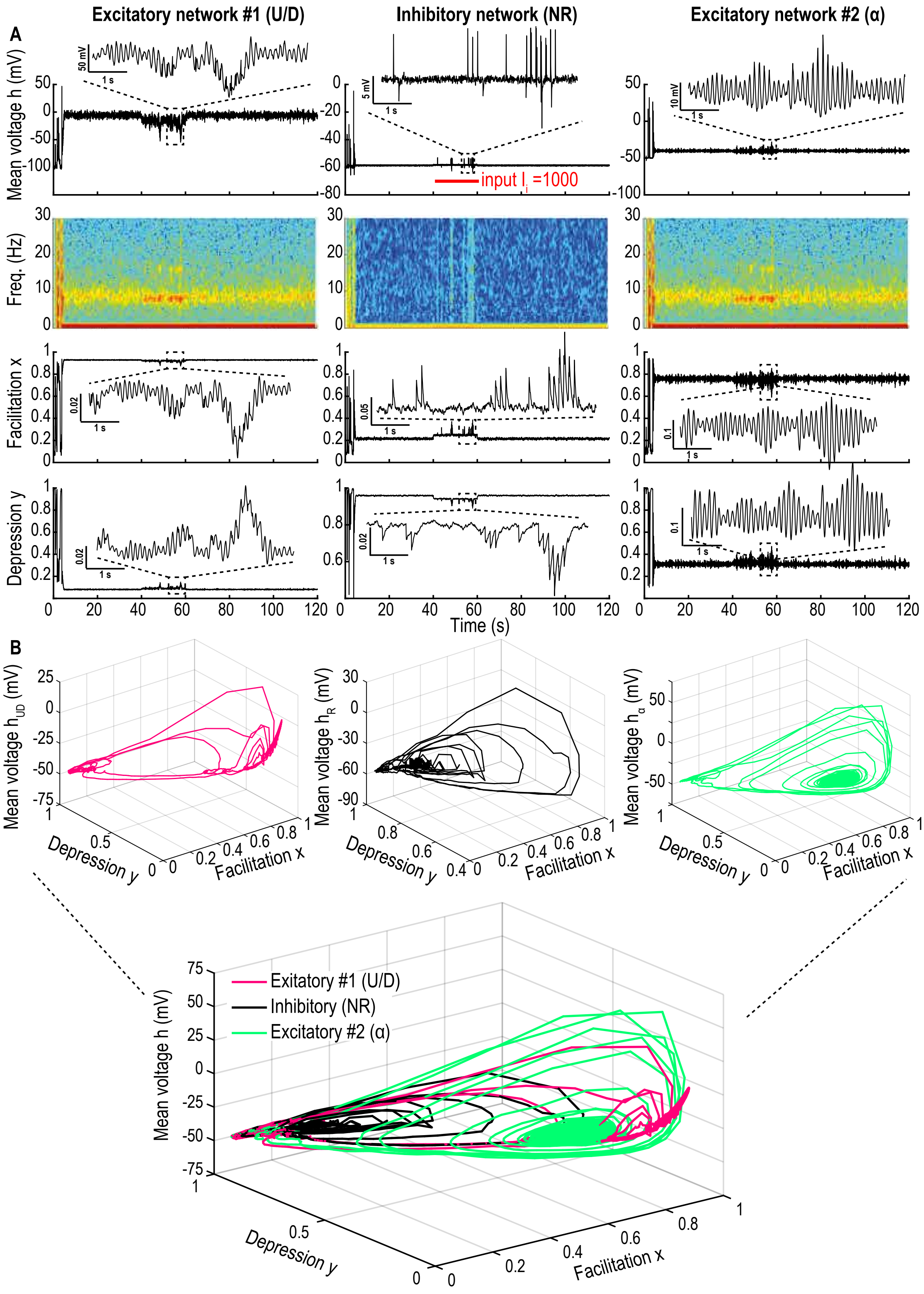


Figure S5: **Contribution of the three components of model (3) for a step input. A.** Time-series of mean voltage $h$, spectrogram, facilitation $x$ and depression $y$ of system 3 (120s simulations) for the excitatory network with AHP ($U/D$, left: $\tau = 0.005$s, $\tau_f = 0.06$s,$\tau_r = 0.12$s), the inhibitory network (NR, center) and the excitatory network without AHP ($\alpha$, right: $\tau = 0.005$s, $\tau_f = 0.06$s,$\tau_r = 0.12$s) with a step input $I_i = 10\cancel{6}0$ at 40-60s on the inhibitory network (red line). **B.** Trajectories in the $h - x - y$ phase space of each component ($U/D$, pink, left, NR black, center and $\alpha$, green, right).

### 2 Supplementary methods

#### 2.1 Fragmentation analysis of an oscillatory band

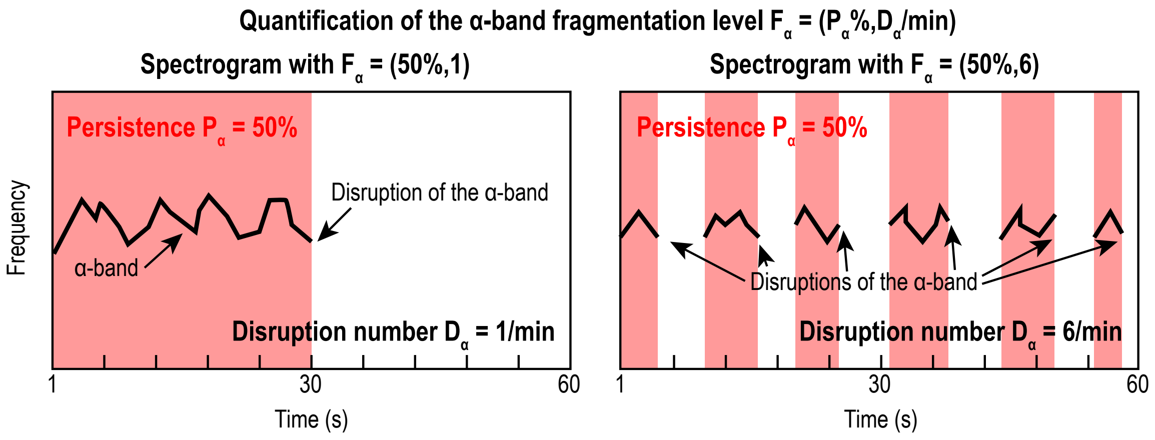


Figure S6: **Schematic of the fragmentation analysis using the spectrogram**

#### 2.2 Mathematical analysis of the phase-space associated with the mean-field depression-facilitation model

We shall now describe the phase-space of the dynamical system (1) with and without AHP. In a first subsection we describe the three critical points (two attractors and a saddle-point) and the linearized dynamics around each point and in a second subsection we describe the numerical method used to obtain the shape of the separatrix delimiting the basins of attraction of each attractor.

##### 2.2.1 Description of the three critical points of the phase-space of network model (1)

The phase-space of the deterministic system (1) contains three critical points that we shall analyze now.

###### Down state attractor point $A_{Down}$

The basin of attraction of the critical point $A_{Down} = (0, X, 1)$ (fig. S7A and S8A, purple) defines the Down state region. The Jacobian at this point is

$$J_{A_{Down}} = \begin{pmatrix} \dfrac{-1 + JX}{\tau} & 0 & 0 \\ K(1-X) & -\dfrac{1}{\tau_f} & 0 \\ LX & 0 & -\dfrac{1}{\tau_r} \end{pmatrix}. \tag{S1}$$

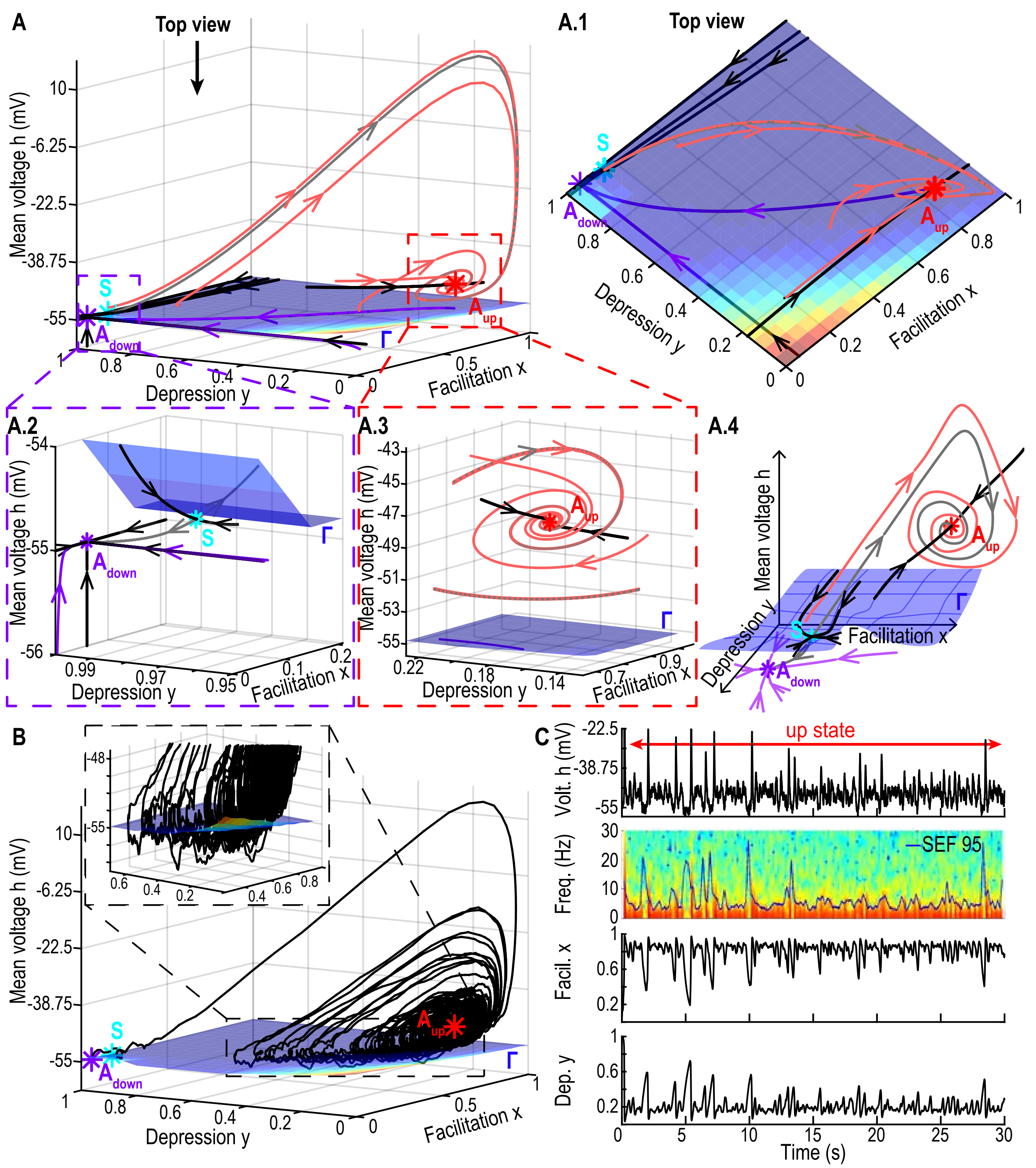


Figure S7: **Phase-space of system (1) without AHP. A.** 3D phase-space of the system with the two attractors $A_{Down}$ (purple, resp. $A_{Up}$, red) and saddle-point $S$ (cyan) with its 2-dimensional stable manifold Γ (blue surface) which defines the separatrix. Stable trajectories (black curves) and unstable manifold of $S$ (grey) and deterministic trajectories starting below (purple, resp. above light red) Γ falling to $A_{Down}$ (resp. $A_{Up}$). Top view (A.1), inset around $A_{Down}$ and $S$ (A.2), inset around $A_{Up}$ where deterministic trajectories oscillate at their eigenfrequency $\omega_{Up}$ (light red, A.3), schematic summary of the entire phase-space (A.4). **B.** Stochastic trajectory lasting $T = 30s$ with $\sigma = 10$ starting at $A_{Down}$ and oscillating around $A_{Up}$. **C.** $(h, x, y)$-time series of a stochastic trajectory, with the spectrogram of the mean voltage $h$ and SEF95 (blue curve).

The eigenvalues are $\left(\lambda_1^{A_{Down}}, \lambda_2^{A_{Down}}, \lambda_3^{A_{Down}}\right) = \left(\frac{JX-1}{\tau}, -\frac{1}{\tau_f}, -\frac{1}{\tau_r}\right)$. When the connectivity $J$ varies in the range $[5.6, 8.6]$, the attractor $A_{Down}$ is a stable-node since the first eigenvalue is negative as long as $J \leq \frac{1}{X} \approx 16.67$. For $J \in [5.6, 8.6], \tau = 0.025s, \tau_f = 0.3s, \tau_r = 0.5s$ and the parameter values of Table 1, we obtain that $\lambda_1^{A_{Down}} \in [-19, -27]$, $\lambda_2^{A_{Down}} \approx -3.33$ and $\lambda_3^{A_{Down}} \approx -2$. The dynamics at this point is identical for the systems exhibiting AHP or not.

#### Up state attractor $A_{Up}$

The second critical point (fig. S7A and S8A, red) is obtained by solving

$$\begin{aligned} x_{Up} &= \frac{\tau_f K(J+1) + LX\tau_r + \sqrt{\Delta}}{2(J\tau_f K + L\tau_r)} \\ y_{Up} &= \frac{1}{Jx_{Up}} \\ h_{Up} &= T + T_0 + \frac{x_{Up} - X}{\tau_f K(1 - x_{Up})}, \end{aligned} \tag{S2}$$

where

$$\Delta = (\tau_f K(J+1) + LX\tau_r)^2 - 4(J\tau_f K + L\tau_r)\tau_f K. \tag{S3}$$

The dynamics around this point depends on whether the system exhibits AHP or not, we will now describe these two cases.

1. **Neuronal network without AHP:** For that system, the resting membrane potential $T_0$ and the recovery timescale $\tau_0$ of the mean voltage $h$ are constant in the entire phase-space. The numerical range of values for the position of the critical point $A_{Up}$ for $J \in [5.6, 8.6], \tau = 0.01s, \tau_f = 0.2s, \tau_r = 0.12s$ and parameters values from Table 1 is $A_{Up} = (h_{A_{Up}} \in [73.15, 124.59]$, $x_{A_{Up}} \in [0.83, 0.89]$, $y_{A_{Up}} \in [0.22, 0.13])$. The Jacobian at this point is

$$J_{A_{Up}} = \begin{pmatrix} 0 & \dfrac{Jy_{Up}(h_{Up} - T - T_0)^+}{\tau_0} & \dfrac{Jx_{Up}(h_{Up} - T - T_0)^+}{\tau_0} \\ K(1 - x_{Up}) & -\dfrac{1}{\tau_f} - K(h_{Up} - T - T_0)^+ & 0 \\ -\dfrac{L}{J} & -Ly_{1,2}(h_{Up} - T - T_0)^+ & -\dfrac{1}{\tau_r} - Lx_{Up}(h_{Up} - T - T_0)^+. \end{pmatrix} \tag{S4}$$

With the present parameters, $J_{A_{Up}}$ has one real negative and two complex conjugate eigenvalues with negative real part thus $A_{Up}$ is a stable-focus: $\lambda_1^{A_{Up}} \in [-55.71, -79.40]$ for the real eigenvalue and the two complex conjugate eigenvalues are

$$\lambda_{2,3}^{A_{Up}} \in [-6.16, -14.73] \pm i[36.78, 51.87].$$

2. **Neuronal network exhibiting AHP:** the Up state attractor $A_{Up}$ is situated in the subspace of medium dynamics with hyperpolarization $\Omega_{mAHP}$ (fig. S8A-B, orange) where $T_0 = T_{AHP} =$

$-30$ and $\tau_0 = \tau_{m,AHP} \in [0.06, 0.3]$s. For $J \in [5.6, 8.6], \tau = 0.025s, \tau_f = 0.3s, \tau_r = 0.5s$ the position of $A_{Up}$ is now $A_{Up} = \left(h_{A_{Up}} \in [-0.74, 19.84], x_{A_{Up}} \in [0.83, 0.89], y_{A_{Up}} \in [0.22, 0.13]\right)$. Here, the eigenvalues of $J_{A_{Up}}$ are real and negative, thus for the system with AHP $A_{Up}$ is a stable-node. The numerical values are now $\lambda_1^{A_{Up}} \in [-34.01, -43.96]$, $\lambda_2^{A_{Up}} \in [-11.67, -18.94]$ and $\lambda_3^{A_{Up}} \in [-3.96, -3.65]$.

##### Saddle-point $S$

The third critical point $S$ (fig. S7A and S8A, cyan) is solution of equations

$$\begin{aligned} x_S &= \frac{\tau_f K(J+1) + LX\tau_r - \sqrt{\Delta}}{2(J\tau_f K + L\tau_r)} \\ y_S &= \frac{1}{Jx_S} \\ h_S &= T + T_0 + \frac{x_S - X}{\tau_f K(1 - x_S)}, \end{aligned} \tag{S5}$$

for $J \in [5.6, 8.6], \tau = 0.01s, \tau_f = 0.2s, \tau_r = 0.12s$ and the parameters are presented in Table 1, we get $A_S = (h_S \in [2.52, 1.08], x_S \in [0.18, 0.12], y_S \in [0.97, 0.99])$. The Jacobian at $S$ does not depend on whether the system exhibits AHP or not and it has one real positive and two real negative eigenvalues, it is thus a saddle-node with an unstable manifold of dimension one and a stable manifold of dimension two. With the present parameters, we obtain $\lambda_1^S \in [-28.80, -25.03]$, $\lambda_2^S \in [18.96, 16.08]$ and $\lambda_3^S \in [-4.89, -4.97]$. Finally, the stable two-dimensional manifold $\Gamma$ defines the separatrix between the basins on attraction of Down $A_{Down}$ and Up $A_{Up}$ states.

##### 2.2.2 Numerical construction of the separatrix

To represent the stable manifold $\Gamma$ of the saddle-point $S$, we use the following algorithm based on numerical approximations (figs. S7A-B and S8A-B, blue surface). Since $\Gamma$ defines the separatrix between the two basins of attraction for the attractors $A_{Down}$ and $A_{Up}$, we ran simulations of the noiseless dynamics for $\sigma = 0$ of system (1) with the initial condition sampling the entire phase space. We used grid points $(h_i, x_i, y_i) \in [-35, 500] \times [0, 1], \times [0, 1])$ with $\delta_h = 1, \delta_x = \delta_y = 0.05$. Each initial point was then attributed to the basin of attraction of the attractor at which the corresponding trajectory ended. The separatrix $\Gamma$ is defined as the border between the set of initial points falling into the basin of $A_{Down}$ and those falling into the basin of $A_{Up}$.
This separatrix does not define a bounded domain for neither attractor but rather separates the entire phase-space in two subdomains, one above $\Gamma$ leading to the Up state and the other one below $\Gamma$ to the Down state .

#### 2.3 Segmentation of the time-series to detect Up and Down states

To determine whether the neuronal population is in an Up or a Down state, we segmented the simulated time-series according to the following criteria:

- the Up states are defined in the subspace $\{x \geq x_{Up} = 0.5 \& h \leq h_{Up} = 0.175 h_{max}\}$,

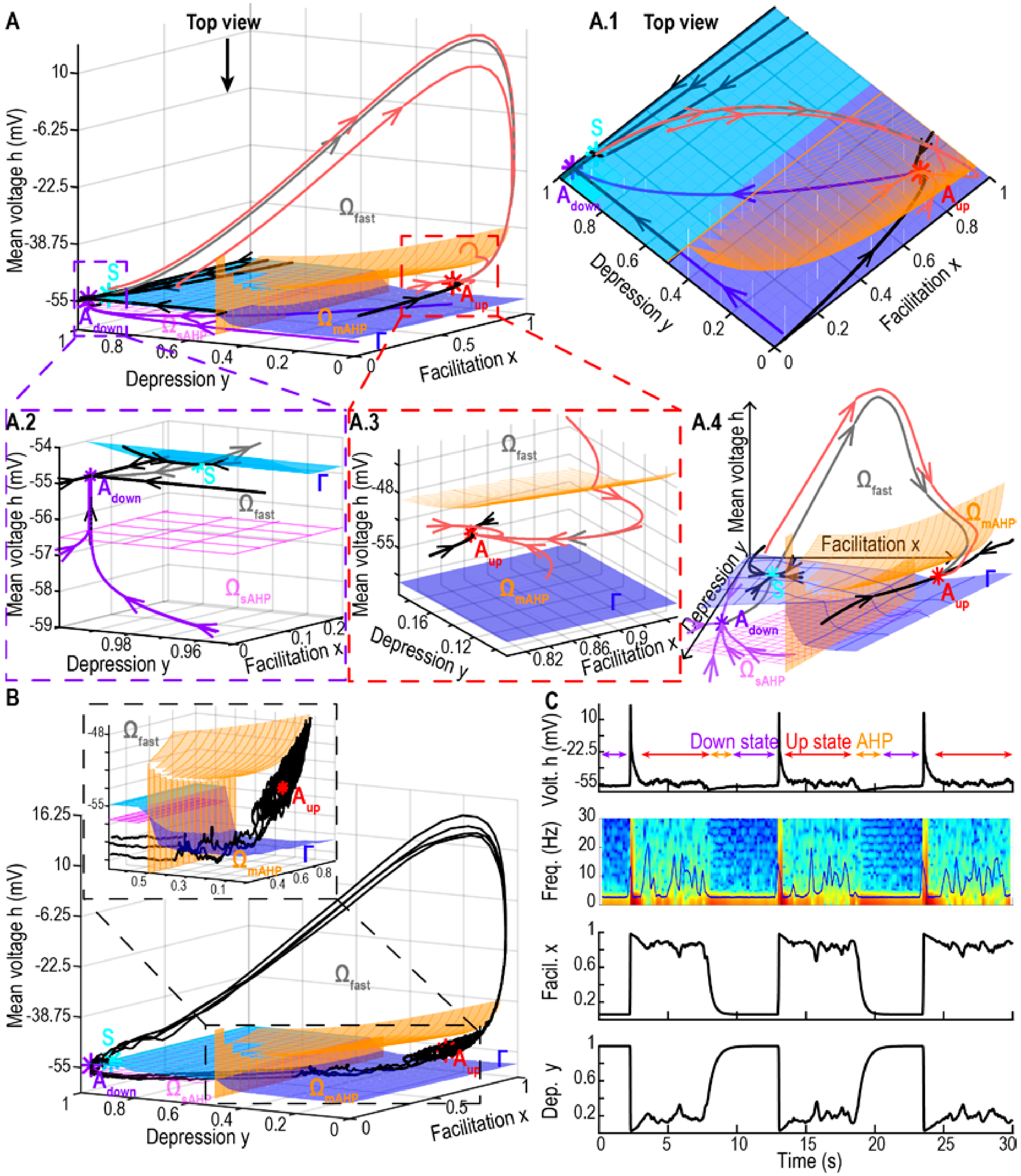

Figure S8: **Phase-space of system (1) with AHP. A.** 3D phase-space of the system with the two attractor points $A_{Down}$ (purple), $A_{Up}$ (red) and the saddle-point $S$ (cyan) with its 2-dimensional stable manifold $\Gamma$ (blue surface) which defines the separatrix. Stable trajectories (black curves) and unstable manifold of $S$ (grey) and deterministic trajectories starting below (purple, resp. above light red) $\Gamma$ falling to $A_{Down}$ (resp. $A_{Up}$). The phase-space is separated into 3 subspaces defining the different dynamics: fast $\Omega_{fast}$ (above pink and orange meshes), medium $\Omega_{mAHP}$ (below the orange mesh) and slow $\Omega_{sAHP}$ (below the pink mesh). Top view (A.1), inset around $A_{Down}$ and $S$ (A.2), inset around $A_{Up}$ (A.3), schematic summary of the entire phase-space (A.4). **B.** Stochastic trajectory lasting $T = 30s$ with $\sigma = 10$ starting at $A_{Down}$ and oscillating between $A_{Up}$ and $A_{Down}$. **C.** $(h, x, y)$-time series of a stochastic trajectory, with the spectrogram of the mean voltage $h$ and SEF95 (blue curve).

- the Down states are defined when $\{y \geq y_{Down} = 0.95\}$

We added the threshold on $h$ for the Up state detection because we do not want to count the bursts, defining the transition from Down to Up, as an Up state.
To determine the proportion of time spent in Up vs Down state for one neuronal population with AHP (fig. 3C-D, main text), we ran simulations of system (1) with AHP for $N = 100$ trajectories of duration $T = 600s$ with $J \in \{5.6, 6.6, 7.6\}$ and $\sigma = 14$.
Similarly, for the model (2) with two populations (fig. 4B-D, main text), we segmented the time-series of the excitatory population for $N = 100$ trajectories of duration $T = 600s$.
Finally for the three population network (3), we segmented the time-series of the excitatory network $\alpha$ without AHP ($N = 100$ trajectories of duration $T = 300s$).

#### 2.4 Numerical methods

All simulations were run in Matlab, using Runge-Kutta 4 scheme with a time step $\Delta t = 0.005$s. We also tried $\Delta t = 0.001$s and obtained the same results, thus ensuring stability.
